# Shifted distribution baselines: neglecting long-term biodiversity records risks overlooking potentially suitable habitat for conservation management

**DOI:** 10.1101/565929

**Authors:** Sophie Monsarrat, Peter Novellie, Ian Rushworth, Graham Kerley

## Abstract

Setting appropriate conservation measures to halt the loss of biodiversity requires a good understanding of species’ habitat requirements and potential distribution. Recent (past few decades) ecological data are typically used to estimate and understand species’ ecological niche. However, historical local extinctions may have truncated species-environment relationships, resulting in a biased perception of species’ habitat preferences. This may result in incorrect assessments of the area potentially available for their conservation. Incorporating long-term (centuries-old) occurrence records with recent records may provide better information on species-environment relationships and improve the modeling and understanding of habitat suitability. We test whether neglecting long-term occurrence records leads to an underestimation of species’ historical niche and potential distribution and identify which species are more vulnerable to this effect. We compare outputs of species distribution models and niche hypervolumes built using recent records only with those built using both recent and long-term (post-1500) records, for a set of 34 large mammal species in South Africa. We find that, while using recent records only is adequate for some species, adding historical records in the analyses impacts estimates of the niche and habitat suitability for fourteen species (41%) in our dataset, and that this effect is significantly higher for carnivores. These results show that neglecting long-term biodiversity records in spatial analyses risks misunderstanding, and generally underestimating, species’ niche, which in turn may lead to ill-informed management decisions, with significant implications for the effectiveness of conservation efforts.

## INTRODUCTION

To avert the ongoing human-induced biodiversity decline, scientists have recently called for conservation efforts to be intensified, including through increased habitat protection and restoration [1]. Data on species’ distribution patterns and species assemblages are key to identify candidate areas for conservation [2]. However, distribution patterns have been drastically modified by humans, notably through global extinctions and regional to local extirpations [3,4], and thus contemporary patterns do not necessarily reflect species’ natural distribution and habitat preferences. Analyses of species distributions that tend to ignore these modifications will likely result in a biased understanding of species’ biogeography and ecological requirements and lead to misleading perceptions of the options available for conservation [5,6]. This phenomenon of spatially shifted baseline poses clear challenges for conservation and management. By providing information on species’ historic rather than current-day relictual distribution, long-term biodiversity data have the potential to improve our understanding of the biogeography of species and participate in setting appropriate spatial and ecological baselines for environmental conservation and restoration [7].

Mammals are one of the most studied taxa, and their current distribution patterns are well known [8]. Historic and prehistoric human-driven global and local extinctions have however caused a strong deviation between current and pre-anthropogenic impact diversity patterns, in particular for large terrestrial mammals [4]. In South Africa, habitat loss, competition with livestock and direct exploitation, in particular following European colonization, have resulted in the global extinction of one mammal species [9], and the collapse of large mammal diversity in large parts of the country [10,11]. To halt this decline and restore populations, conservation efforts have focused on establishing protected areas and actively managing large mammal populations through reinforcement - to increase population viability - and reintroductions - to re-establish populations within species’ historical range [12]. Defining species’ historical distributions and suitable habitat is thus a critical aspect for conservation planning in South Africa [13], as it is for most restoration attempts elsewhere [14]. Among large mammal species, those identified as threatened by the IUCN Red List are a high priority for conservation [15] and large carnivores have an important ecological role and have undergone considerable historic range contractions [16], making them a major focus of conservation and rewilding efforts [17]. It is thus critical to understand the extent to which historical data are needed to inform wildlife conservation and management, for threatened species and large carnivores in particular.

Habitat suitability models (HSMs) [18,19] and *n*-dimensional hypervolumes [20] are two widely-used tools that relate species’ occurrences to environmental variables in order to, respectively, map species’ potential distribution in geographical space and characterize species’ niche in the environmental space. They have notably been used in a conservation and management contexts to improve our knowledge of species ranges, support management plans for species’ recovery, prioritize areas for biodiversity protection and predict changes in suitable habitat in response to human impacts [21–23]. HSMs and hypervolume approaches rely on the assumption that the observed geographical distribution of a species reflects its ecological requirements, making them highly contingent on the quality of the occurrence records[19,23]. Range contraction that have affected the array of conditions that the species occupy risk truncating species–habitat relationships [24], thus hindering our ability to estimate species niches and predict the distribution of suitable habitat [6]. Failing to consider past local extinction events may thus misguide conservation efforts by overlooking potentially suitable sites for reintroduction or restrict protection to suboptimal habitats [5]. Despite providing useful information on the historic distribution of species, historical written records and museum specimens have long been overlooked in habitat suitability modeling approaches, being perceived as untrustworthy for their intrinsic biases and limitations [25] (but see [26–28]). The development of methods to address sampling biases in HSMs [29–32] however, provides an avenue for more confident incorporation of these records in spatial modeling analyses, and hence in conservation interventions.

Here we investigate how long-term biodiversity records can contribute to setting appropriate baselines for species’ distribution. We test the hypothesis that neglecting historical data in niche quantification and HSM approaches leads to biased perceptions of species’ historic niche and suitable habitat distribution, and the patterns of potential species richness at the regional level. We focus on large terrestrial mammals in South Africa, for which we have access to a unique dataset of long-term occurrence records spanning the last five centuries, as well as recent (post-1950) occurrence records, for a community of 34 mammal species.

## METHODS

### Overview of the approach

We considered two datasets of occurrence - recent records (post-1950, RECENT) and recent + long-term records (post-1500, TOTAL) - to quantify the effect of neglecting long-term occurrence data. We compared results obtained from these two datasets in three different approaches, two that are species based: 1) estimation of the climatic niche in environmental space using n-dimensional hypervolumes [23] and 2) prediction of suitable habitat in the geographic space using habitat suitability models (HSMs) and one at the community level, namely prediction of the distribution of potential species richness using stacked-HSMs. For the species-level approaches, we used two indices that summarize the cost of neglecting long-term biodiversity data and tested how the combination of these indices relate to species’ conservation status and diet. For the community-level approach, we investigated spatial differences in predicted potential species richness, notably by comparing predictions between different South African bioregions.

### Species data

The general study area, hereafter referred to as South Africa, covers the countries of South Africa, Lesotho and eSwatini (former Swaziland). We considered all extant South African large (> 20 kg) terrestrial mammals, except for species with fewer than 25 long-term observations in the dataset. In total, 34 species were included: 23 from the order Artiodactyla, 6 Carnivora, 4 Perissodactyla and 1 Proboscidea.

#### Theoretically accessible areas

Barve et al. [33] outline the concept of the theoretically accessible area (the area that is climatically suitable and has been accessible to the species via dispersal over relevant periods of time), and show that restricting a model’s training and validation areas to this theoretically accessible area greatly improves HSM performance and provides more accurate predictions of species richness and community composition [33,34]. As an approach to estimating the theoretically accessible area for each species, we identified the bioregions in which the species are known to have occurred historically, based on information on their ecology and interpretation of historical occurrences, and built a polygon using the boundaries of these bioregions. We defined the accessible area for each species as a buffer of 20 km around this polygon, to include ecotone regions where the species could disperse. This option is suggested by Barve et al. [33] to be the most operational compared to more intricate alternatives. We acquired spatial information on bioregions from the 2012 Vegetation Map of South Africa, Lesotho and Swaziland [35].

#### Modern and historical occurrence records

The long-term occurrence dataset used in this study covers the period 1500 to 1950 and includes records extracted from the historical literature, museum specimens and fossil records. For historical records and museum specimens, we used the database presented in Boshoff et al [10], completed with records from the KwaZulu-Natal, eSwatini and the rest of South Africa, using the same approach and criteria defined in Boshoff et al. [10], so that the dataset covers all of South Africa. The reliability of these records in terms of identification and locality is discussed in Boshoff and Kerley [36] and their spatial, environmental and taxonomic biases in Monsarrat et al. [29] and Monsarrat and Kerley [37].

Recent fossil records were obtained from Avery [38], a comprehensive compilation of information on the taxonomy and distribution in time and space of all currently recognized South African fossil mammal. We recovered radiocarbon dating for these record from primary sources and kept only those that were deposited in the period from 1500 to today.

Modern (post-1950) occurrence records were provided by the South African National Biodiversity Institute and the Endangered Wildlife Trust, who consolidated and centralized a total of over 460,000 geo-referenced unique occurrence records for South African mammals, from 59 different contributors (governmental institutions, research institutions, non-governmental organizations, the private sector and citizen science projects) [39]. This database formed the basis of the 2016 “Red List of Mammals of South Africa, Lesotho and Swaziland” and as part of this process, data were vetted and underwent several rounds of data cleaning to check accuracy. These data are spatially biased, with the highest densities of records typically found in protected areas [39], artificially increasing spatial auto-correlation of occurrences. This in turn may affect the performance of habitat suitability models built with these data [40]. To reduce the effect of sampling bias and spatial clustering on model performance, we subsampled the modern occurrence dataset using spatial thinning of the data (no occurrence records closer than 0.1 degree), as recommended by Boria et al. [41].

We considered all occurrence records located outside of a species’ theoretically accessible area to be extralimital and we excluded them from the analyses. Modern extralimital records often correspond to introductions of individuals or populations outside of their historic range, often in suboptimal habitat, and are not informative of the habitat preferences of the species [42]. We however acknowledge that, by using bioregions as the filter for modern records, we may include some records that are outside the historic range, this being due to the relatively unique situation in SA of game translocations for commercial purposes [42].

In total, we analyzed 15,315 recent (post-spatial thinning, range 55-1,274) and 5,446 historical (range 25-501) records for the 34 species of large terrestrial mammals, covering a total area of ca. 1,270,000 km^2^ for these three nations.

## Environmental data

We considered six bioclimatic variables derived from BioClim [43]: mean annual temperature (BIO1) and annual precipitation (BIO12), describing the average climatic conditions, temperature seasonality (BIO4) and precipitation seasonality (BIO15), describing climatic seasonality and maximum temperature of the warmer month (BIO5) and precipitation of the warmest quarter (BIO18), describing extreme climatic conditions. We also considered topography (TOPO), using altitude data from the ASTER Global Digital Elevation Model (ASTGTM) on https://lpdaac.usgs.gov [44]. These variables were chosen, because they were biologically meaningful to predict large mammal species richness in South Africa [45] and because they potentially represent environmental characteristics that limit species’ distributions. All environmental variables were estimated at a 0.1 x 0.1 degree resolution, using the *raster* package [46] in R 3.5.1 [47].

## Hypervolume analysis

The n-dimensional hypervolume was originally proposed by Hutchinson [48] to describe the fundamental niche of a species, i.e. the environmental space where the species can exist indefinitely. In the modern understanding of the hypervolume function, a set of *n* variables that represent biologically important and independent axes are identified and the hypervolume is defined by a set of points within this n-dimensional space that reflects suitable values of the variables for the species’ persistence [23]. Here, we consider five environmental axes: BIO1, BIO4, BIO12, BIO15 and TOPO, rescaled to a common and comparable scale before the analyses. We used the Gaussian kernel density estimation with the Silverman bandwidth estimator method in the *hypervolume* package [20] in R 3.5.1 [47]. The bandwidth was estimated from the RECENT dataset with the Silverman estimator and the same value was used for the TOTAL dataset, to allow direct comparison.

The volume of the hypervolume is approximately linearly proportional to the number of observations in the dataset [20]. To ensure results are insensitive to sample size, we randomly subsampled the TOTAL dataset to have the same number of records as the RECENT dataset. We repeated the process ten times and used averaged hypervolume measures of these ten repetitions in the statistical analyses.

## Habitat Suitability Modeling

### Background data

Because the species occurrence records are highly biased spatially [29,39], we addressed the potential effect of sampling bias in the models. To do so, we produced background data with similar geographical bias as the RECENT and TOTAL occurrence datasets, following [30]. We first created a sampling effort raster using a two-dimensional kernel density estimation applied on the occurrence dataset. Background data were then created by sampling without replacement within this raster grid, where the probability of a cell being sampled was proportional to the sampling density values (weighted target group approach, following Sanín and Anderson [49]). We selected the same number of background points as the number of occurrence records, so as to achieve a prevalence of 50%, as advised by Liu et al. [50].

### Ensemble modeling

We created ensemble HSM [51] for each species by assembling five statistical methods (GAM, MAXENT, MARS, RF and GBM) to account for inter-model variability, using the *ssdm* package [52]. We ran ten repetitions for each of the algorithms and produced an average of the models’ outputs, weighting each model according to its predictive ability. We measured predictive ability with a cross-validation approach, by using a random 70% of the data for calibration of the models (keeping the prevalence constant) and testing their predictive ability on the remainder of the dataset using the True Skill Statistic (TSS) [53]. We repeated this approach ten times for each model and used an average of the predictive accuracy measure. In total, for each species and each dataset, we ran 500 models using five different statistical models, ten repetitions of each algorithm, and ten repetitions of the random-splitting strategy. The outputs of these models are maps of predicted habitat suitability over the study area that provide hypotheses for the potential distribution of species for both datasets. We identified areas where the predicted habitat suitability differs between the RECENT and TOTAL datasets by subtracting the predicted values obtained from the RECENT model to those obtained with the TOTAL model in each cell within the study area. Areas with positive (negative) values are where we underestimate (overestimate) habitat suitability when considering only recent records.

### Species richness

Stacked-species distribution models (SSDM) combine multiple individual HSMs to produce a community-level model and predictive maps of potential species richness [54]. We used the *ssdm* package [52] to compute maps of local species richness by summing the probabilities from continuous habitat suitability maps provided by the ensemble HSMs, a method that performs better than stacking methods based on thresholding site-level occurrence probabilities [55]. To highlight areas where the potential species diversity is under-or over-estimated because of neglecting long-term occurrence records in the models, we subtracted the map of species richness produced with the RECENT dataset to the one produced with the TOTAL dataset. We also compared the mean difference in predicted species richness for each bioregion of South Africa.

### Statistical analyses

For each species, we considered two indices to summarize the effects of neglecting long-term records on the estimation of climatic niche and habitat suitability: 1) the niche dissimilarity in environmental space (N_dis_) [20] and 2) the dissimilarity in predicted habitat suitability in the geographical space (PRED_dis_) [56]; (see Table 1 for a definition of these indices). For each index, higher values indicate higher disparity between the results obtained with the RECENT and the TOTAL dataset.

**Table 1.**
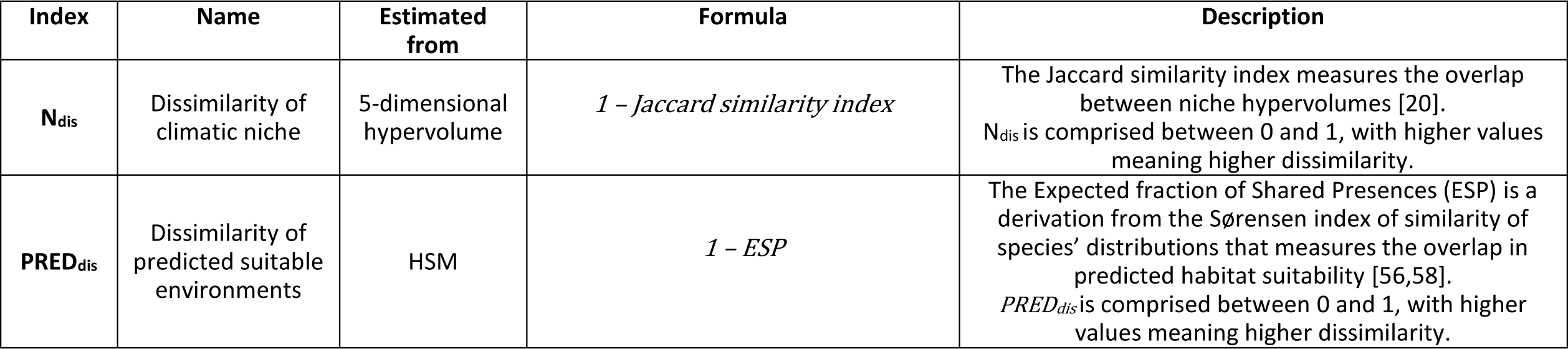
Description of the two indices used in the PCA analysis to quantify the effect of neglecting historical records on the estimation of climatic niche and suitable habitat.

We rescaled all indices by subtracting the mean and dividing by the standard deviation so that they are comparable and conducted a Principal Component Analysis (PCA) to convert these indices into a one-dimension variable (the first principal component PC1), quantifying the effect of neglecting historical records. We ran a two-way ANOVA with Type II errors to test for differences in PC1 between conservation status (threatened vs non-threatened) and broad diet guilds (herbivores vs carnivores). The conservation status was defined from the IUCN Red List categories [57], where species listed as vulnerable, endangered or critically endangered were considered “threatened”, and “non-threatened” otherwise. We used a linear model to test how the change in mean predicted habitat suitability (ΔPRED, calculated as the proportion difference in mean predicted habitat suitability over the study area when it is estimated from the TOTAL dataset, compared to the RECENT dataset) varies with PC1 values. We also estimated the difference in the ability of HSMs to predict all the known occurrences for the species (ΔB), by measuring the proportional increase (or decrease) in the continuous Boyce index, a threshold-independent evaluator of the ability of HSMs to predict species presences [54], when it is estimated from the TOTAL dataset, compared to the RECENT dataset.

## RESULTS

The ensemble modeling approach yielded very good agreement between the different modelling methods, as indicated by low standard deviation around the predicted habitat suitability values (Supplementary Information 2). For 18 out of 34 species, the inclusion of historical records improved the ability of the model to predict all known occurrences of the species (ΔB>0). The highest improvement in predictive ability was for the Black rhinoceros, Bontebok and African Elephant (ΔB equal to 25%, 16% and 11%, respectively). In contrast, eleven species showed a decrease in predictive ability when historical data are included in the model, with the gemsbok and blue wildebeest showing the strongest decrease (ΔB equal to −10% and −8%, respectively). The dissimilarity in climatic niche N_dis_ calculated from the 5-dimensional hypervolume ranges from 0.09 to 0.52 (mean=0.21 ± 0.11 SD), with N_dis_ > 0.40 for the African elephant, lion and African wild dog. The dissimilarity in predicted habitat suitability PRED_dis_ ranges from 0.41 to 0.71 (mean=0.57 ± 0.07 SD), with PRED_dis_ > 0.70 for the African wild dog, lion and spotted hyaena (Table S1 of Supplementary Information).

Overall, by combining N_dis_ and PRED_dis_ in a PCA, fourteen species (41% of our dataset) come out as impacted by neglecting historical records (PC1 < 0, with PC1 explaining 79% of the variance), with five species identified as the top most impacted: the lion, African wild dog, African elephant, spotted hyaena and hippopotamus (Fig 1A). Of these five species, four are listed as threatened on the IUCN Red List. Three are carnivores and the other two are megaherbivores (body mass >1,000 kg). PC1 values were significantly lower for carnivores compared to herbivores (two-way ANOVA Type II, F(1,34)= 5.35, MSE = 6.60, p=0.026), and marginally lower for threatened compared to non-threatened species (two-way ANOVA Type II, F F(1,34)=2.26, MSE = 2.78, p=0.143).

**Figure 1.**
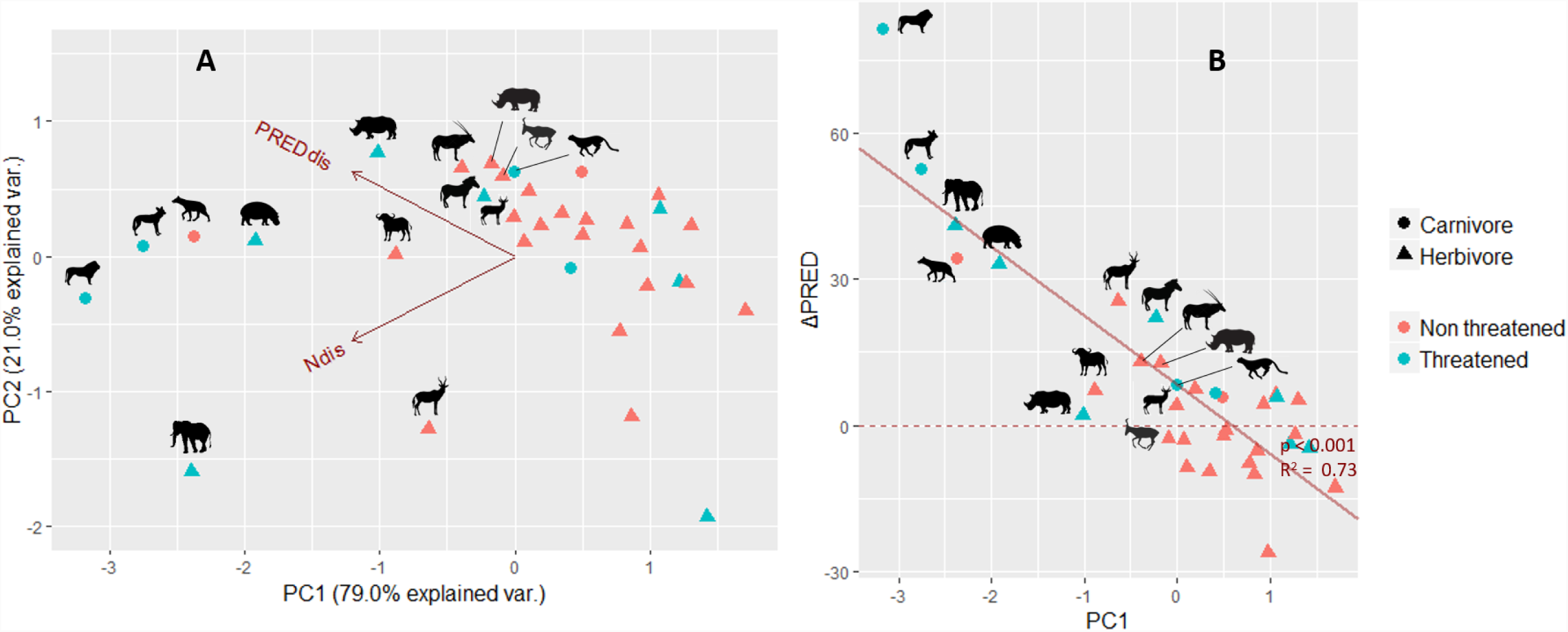
Effects of incorporating historical records on the estimation of species climatic niche and predictions of habitat suitability. A) Principal Component Analysis (PCA) of the two indices used to measure discrepancy between estimations of climatic niche (N_dis_) and habitat suitability (PRED_dis_) with the RECENT and TOTAL datasets, for the 34 species of large mammals considered. Higher values of indices (lower values of PC1) indicate a higher discrepancy. We highlighted (silhouettes) 14 species with negative PC1 values, most affected by neglecting historical records. We differentiate carnivores vs herbivores and threatened vs non-threatened species. The differences between these groups along the first principal component (PC1) are significant for the former (p=0.026) and marginal for the latter (p=0.143). B) Plot showing the negative relationship between values of PC1 and the proportion difference in mean predicted habitat suitability over the study area when it is estimated from the TOTAL dataset, compared to the RECENT dataset (ΔPRED). Positive values of ΔPRED mean that the habitat suitability is underestimated without using historical records. Species that are most affected by neglecting historical data have increased mean predicted habitat suitability when historical records are included in HSMs. See Table S1 in Supplementary Information for a key of silhouettes.

We found a significant inverse linear relationship between PC1 and the change in mean predicted habitat suitability over the study area ΔPRED (p<0.001, R_^2^_=0.73) (Fig 1B), i.e. species that are most affected by neglecting historical data have higher mean predicted habitat suitability over their study area when historical records are included in HSMs. The lion, African wild dog, elephant, spotted hyaena and hippopotamus show the largest increase in mean predicted habitat suitability when historical records are included (ΔPred equal to 81%, 53%, 41%, 34% and 33%, respectively).

This results in differences in predicted potential species richness at the community level (Fig 2A) and in the geographic distribution of predicted habitat suitability at the species level (see maps on Fig 2B for the five most impacted species and Supplementary Information for maps of all 34 species). Differences in predicted potential species richness are higher for the Albany Thicket, Fynbos and Savanna Lowveld biomes (Fig 3). The Nama-Karoo has on average very similar predicted species richness with the RECENT or TOTAL dataset, whereas the potential species richness tends to be overestimated in Arid Savanna.

**Figure 2.**
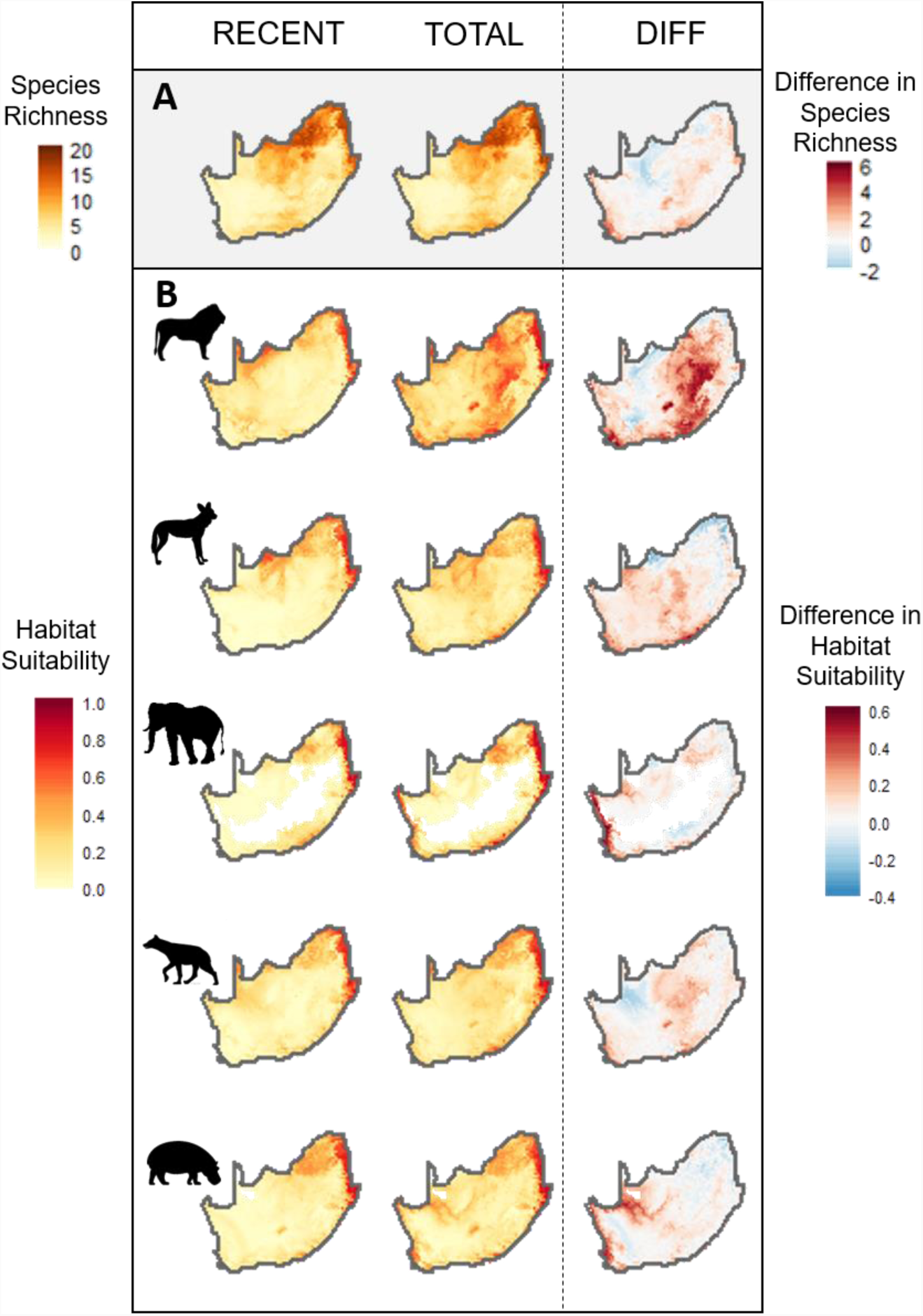
Effect of incorporating historical records in the spatial prediction of A) species richness and B) habitat suitability for the five species most impacted by neglecting historical records. The first column is the prediction of species richness/habitat suitability obtained from the RECENT dataset (post-1950 records) and the second column is obtained from the TOTAL dataset (historical + recent records). The last column is the difference between column 2 and column 1. We highlight the five species most impacted through neglecting historical records according to their PC1 score: lion, African wild dog, African elephant, spotted hyaena and hippopotamus.

**Figure 3.**
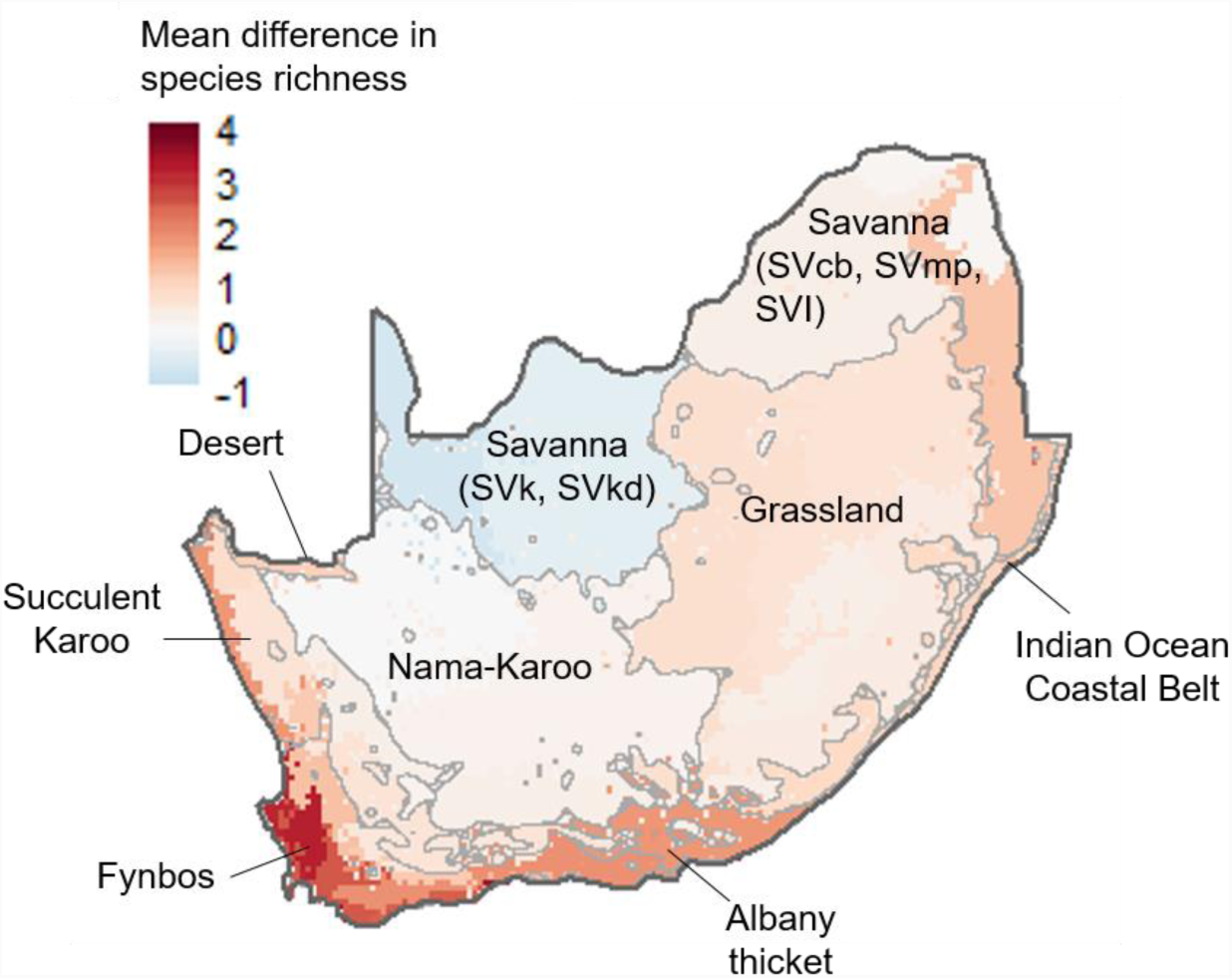
Mean difference in species richness estimated from the TOTAL vs RECENT dataset, calculated for each of the bioregions of South Africa. Darker shades of red indicate that the potential species richness is higher when predicted with the TOTAL dataset than with the RECENT dataset. The main biomes are identified with grey contouring, obtained from simplifying polygons of the 2012 “Vegetation Map of South Africa, Lesotho and Swaziland” [59]. The distinction is made between arid Savannas (SVk: Eastern Kalahari Bushveld Bioregion and SVkd: Kalahari Duneveld Bioregion) and mesic Savannas (SVcb: Central Bushveld Bioregion; SVmp: Mopane bioregion; SVl: Lowveld Bioregion and SVs: Sub-Escarpment Savanna Bioregion) [59].

## DISCUSSION

We show that neglecting long-term records can bias estimates of species climatic niche and suitable habitat and underestimate the potential regional species richness. The implications are more severe for carnivore species, and marginally more so for threatened species, for which appropriate conservation actions and management decisions are the most critical. These results have implications for conservation planning and distribution modelling in general, given that globally most mapping of species’ distributions and habitat use exclude long-term occurrence records, and for the conservation and management of South African mammalian fauna. These findings highlight the importance of considering long-term data in modern ecological analyses and may also provide explanatory insights into limits of conservation approaches when they fail to consider appropriate species distribution baselines. We expand on these points below.

### Species implications

For several species, we observe only a limited effect of including historical records in the analyses. This indicates that modern occurrence records provide a reasonably good coverage of the climatic conditions found in their historic distribution. This possibly reflects that they have been less impacted by past range contractions, that range contraction did not affect the range of environmental conditions occupied by the species or that they have successfully recovered throughout their historic range, whether by natural recolonization or through active reintroductions. This result highlights the success of conservation efforts in South Africa, where many species have been successfully reintroduced throughout their historic range (e.g. black wildebeest, Cape mountain zebra) [60], with some species even introduced outside their native range (e.g. giraffe, impala) [42] (though extralimital records have been excluded from the analyses and thus the consequences of these introductions were not reflected in this study).

In contrast, considering historical data hugely affects estimates of the climatic niche and potential distribution of other species. For these, the geographic distribution of predicted habitat suitability is wider than expected from recent data only and this effect is higher for species of high conservation value. Three of the five most impacted species are carnivores (lion, African wild dog and spotted hyaena) and the two others are megaherbivores (elephant and hippopotamus), all being listed as threatened by the IUCN Red List except the spotted hyaena, which is revealingly listed as near threatened on the Red List of Mammals of South Africa, Swaziland and Lesotho [61]. These species are highly charismatic [62], very sensitive to humans [63], and play important roles in ecosystems[64], thus acting as focal species for management efforts and trophic rewilding initiatives [65]. Analyses based on recent data only will lead to truncated estimates of bioclimatic relationships and underestimations of the extent of suitable areas for conservation. Important suitable areas might be overlooked when selecting appropriate sites for reintroductions and trophic rewilding, and protection efforts might focus on marginal habitat [5,6]. The implications of such missed opportunities on management outcomes and the conservation status of species need to be better understood.

### Community implications

At the community level, neglecting historical records underestimates potential regional species richness, with some areas being more impacted than others. In South Africa, the south-western and western parts of the coastline as well as the central Free State and the Eastern Cape provinces have higher potential richness than expected from recent records only. These areas were highly impacted historically, with the establishment of the Cape Colony by the Dutch in the mid-17^th^ century and the subsequent colonization of the interior, with increased pressures from land-use change and direct hunting [11,66]. In most bioregions, overlooking historical records underestimates the potential species richness, with particularly strong effects in the Fynbos and Albany Thicket biomes. These shifted distribution baselines clearly have implications for our understanding of broader biogeographic patterns and processes. As an example, the underestimate of large mammal species richness in the Fynbos biome illustrated here demonstrates that the role of mammals in this biome, traditionally considered to support a low diversity of large mammals [13,68], needs to be reassessed. In addition to having suffered major biodiversity declines in the past [13,67], these areas are also where conservation efforts are thus most likely to be misguided (but see [13] for conservation planning in the Fynbos), which carries major implications for wildlife management and conservation in South Africa.

Our study area doesn’t cover the full distribution range of some species, meaning that we are only sampling part of the environmental conditions that these species might encounter throughout their range. While this limits the transferability of predictions in space or time, our results remain valid at the regional level because we don’t extrapolate outside the environmental space sampled in the occurrence dataset. South Africa is an exceptional ecoregion, with unique climatic regimes, high species richness and endemism [69,70]. Being at the southern margin of some species’ global distribution, it is a particularly important area for conservation since it may harbor populations with unique local adaptations that will be critical for species’ ability to persist in the face of future climate change [71]. Range contractions that truncate species-climate relationships in this area are thus even more critical for our understanding of species’ niche than those occurring at the center of the range.

### Implications for informing climate change and invasion risk

HSMs are widely used to forecast species range shifts under contemporary climate change [72] and to assess the geographic risk of species’ invasions [73]. In a study investigating the impact of overlooking historical records in HSMs for 36 North American mammal species, Faurby and Araújo found that forecasts of climate change impacts on biodiversity are unlikely to be reliable without acknowledging for past anthropogenic range contraction [74]. Our study provides further evidence that using recent distribution records only can underestimate species’ bioclimatic niche, which in turn is likely to provide biased forecast of species’ response to climate change. Similarly, this truncated understanding of suitable bioclimatic areas will make us vulnerable to underestimating invasion risk. While this latter aspect is of less relevance for large mammals which are less frequently involved in invasions, the principle is of importance when modelling risk areas for known invasive species.

### Setting baselines

Shifting baselines [75] emphasize the need for setting appropriate references when exploring ecological patterns and how these may change, especially for detecting long term processes. We have demonstrated here the occurrence of shifted baselines for the distribution of South African mammals, against which one can assess recent and future shifts in the geographic patterns of this fauna. Such phenomenon can be expected elsewhere, and there is thus a need to study this at a global scale.

This study focuses on Pre-European settlement conditions, which are often held up as a relevant baseline from which to define restoration objectives and quantify success [13,76,77]. For South Africa, we consider the 15_th_ century baseline to be appropriate for identifying species’ niche since the bulk of human-related impact for extant large mammal species occurred after this period [11] (but see [78]). But this might not hold true in other systems, where human impact on mammal megafauna occurred much earlier [79]. The appropriate baseline should be adapted accordingly, to estimate natural diversity patterns and allow the identification of sites that match the biotic and abiotic needs of the focal species.

## CONCLUSION

The recognition that neglecting long-term biodiversity might lead to setting inappropriate spatial baselines is the first step towards a better integration of these data in decision-making for biodiversity conservation and management. Due to the difficulty in collecting historical occurrence records, long-term datasets are not currently available for all taxa or regions. However, with the recent recognition of the value of these datasets for conservation, there is an encouraging development towards assembling long-term biodiversity datasets [e.g. 80–82], including for underrepresented taxa [e.g. 83,84]. The release of global databases of historic distributions [85] is a promising avenue to integrate long-term perspectives in future ecological studies. We join previous calls for international, multidisciplinary effort to compile historical data [86], and urge that, whenever possible, these should be included into conservation and biogeography studies. Unless efforts are made to integrate this historical perspective into biodiversity conservation, shifted distribution baselines risk undermining our efforts to define appropriate protected areas and halt the ongoing biodiversity crisis, as well as appropriately manage biodiversity under global change.

## Supporting information

Supplementary Information 1 - Results table

Supplementary Information 2 - HSMs results and maps

## ACKNOWLEDGMENTS

We thank Margaret Avery and Shaw Badenhorst for assistance regarding the archeological records, Matthew Child and the Endangered Wildlife Trust for providing access to the recent occurrence dataset and Ana Rodrigues, for providing comments on an earlier version of the manuscript.

## SUPPLEMENTARY INFORMATION

*Supplementary Information 1:* Summary table of results for the 34 species of large mammals included in the analyses.

*Supplementary Information 2:* HSMs results and maps for 32 species of large mammals included in the analyses (spatial data for the two species of rhinoceros are not provided for security reasons).

