## Supplementary Information 1 - Results table for "Shifted distribution baselines: neglecting long-term biodiversity records risks overlooking potentially suitable habitat for conservation management"

**Table S1.** Summary of results for the 34 species of large mammals included in the analyses, ordered by increasing values of PC1, i.e. the species higher in the list are those for which the effect of neglecting historical data on the estimation of climatic niche and habitat suitability is stronger. C=Carnivore; H=Herbivore; T=Threatened (i.e. listed as Vulnerable; Endangered or Critically Endangered by the IUCN Red List); NT=Not Threatened (i.e. listed as Least Concern or Near Threatened by the IUCN Red List); AUC=Area under the receiver operating characteristic curve; B=Boyce index; ΔPRED=proportion difference in mean predicted habitat suitability; $N_{dis}$= Environmental niche dissimilarity index; $Pred_{dis}$= predicted habitat suitability dissimilarity index; PC1= first axis of the principal component analysis.

| **Species** | Latin name | Diet | Status | $AUC_{recent}$ | $AUC_{tot}$ | $B_{recent}$ | $B_{tot}$ | ΔPRED | $N_{dis}$ | $PRED_{dis}$ | PC1 |
|---|---|---|---|---|---|---|---|---|---|---|---|
| **Lion** | *Panthera leo* 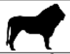 | C | T | 0.83 | 0.76 | 0.93 | 0.97 | 81.48 | 0.48 | 0.71 | -3.17 |
| **African wild dog** | *Lycaon pictus* 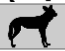 | C | T | 0.84 | 0.78 | 0.81 | 0.90 | 52.68 | 0.41 | 0.71 | -2.75 |
| **African elephant** | *Loxodonta africana* 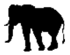 | H | T | 0.86 | 0.86 | 0.94 | 0.97 | 40.94 | 0.52 | 0.61 | -2.39 |
| **Spotted hyaena** | *Crocuta crocuta* 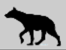 | C | NT | 0.83 | 0.79 | 0.92 | 0.99 | 34.2 | 0.38 | 0.70 | -2.37 |
| **Hippopotamus** | *Hippopotamus amphibius* 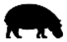 | H | T | 0.83 | 0.82 | 0.92 | 0.99 | 33.16 | 0.35 | 0.67 | -1.91 |
| **Black rhinoceros** | *Diceros bicornis* 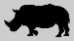 | H | T | 0.68 | 0.73 | 0.94 | 0.98 | 2.03 | 0.23 | 0.66 | -1.01 |
| **African buffalo** | *Syncerus caffer* 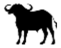 | H | NT | 0.79 | 0.81 | 0.92 | 0.98 | 7.07 | 0.28 | 0.62 | -0.88 |
| **Blesbok** | *Damaliscus pygargus phillipsi* 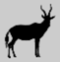 | H | NT | 0.81 | 0.79 | 0.82 | 0.98 | 25.41 | 0.36 | 0.54 | -0.63 |
| **Gemsbok** | *Oryx gazella* 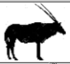 | H | NT | 0.78 | 0.77 | 1.00 | 0.99 | 13.1 | 0.19 | 0.62 | -0.39 |
| **Cape mountain zebra** | *Equus zebra* 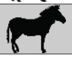 | H | T | 0.73 | 0.77 | 0.99 | 0.94 | 22.12 | 0.19 | 0.61 | -0.23 |
| **White rhinoceros** | *Ceratotherium simum* 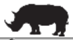 | H | NT | 0.69 | 0.74 | 0.95 | 0.96 | 12.79 | 0.17 | 0.61 | -0.17 |
| **Tsessebe** | *Damaliscus lunatus* 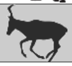 | H | NT | 0.67 | 0.75 | 0.97 | 0.96 | -2.67 | 0.17 | 0.61 | -0.09 |
| **Grey rhebok** | *Pelea capreolus* 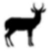 | H | NT | 0.87 | 0.85 | 0.99 | 1.00 | 4.09 | 0.19 | 0.59 | -0.008 |
| **Cheetah** | *Acinonyx jubatus* 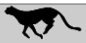 | C | T | 0.82 | 0.83 | 0.99 | 0.97 | 8.24 | 0.16 | 0.60 | -0.001 |
| **Red hartebeest** | *Alcelaphus buselaphus* 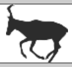 | H | NT | 0.75 | 0.74 | 0.97 | 1.00 | -3.1 | 0.20 | 0.57 | 0.06 |
| **Bushpig** | *Potamochoerus larvatus* 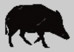 | H | NT | 0.64 | 0.73 | 0.98 | 0.90 | -8.61 | 0.16 | 0.59 | 0.1 |
| **Eland** | *Tragelaphus oryx* 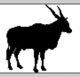 | H | NT | 0.73 | 0.72 | 0.98 | 0.96 | 7.41 | 0.18 | 0.57 | 0.19 |
| **Southern reedbuck** | *Redunca arundinum* 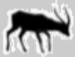 | H | NT | 0.71 | 0.72 | 0.98 | 0.97 | -9.44 | 0.16 | 0.57 | 0.34 |
| **Leopard** | *Panthera pardus* 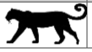 | C | T | 0.85 | 0.85 | 0.99 | 0.97 | 6.55 | 0.18 | 0.55 | 0.4 |

| | | | | | | | | | | | |
|---|---|---|---|---|---|---|---|---|---|---|---|
| **Brown hyaena** | *Parahyaena brunnea* 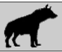 | C | NT | 0.77 | 0.76 | 0.99 | 0.96 | 5.92 | 0.12 | 0.58 | 0.48 |
| **Bushbuck** | *Tragelaphus scriptus* 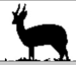 | H | NT | 0.74 | 0.74 | 0.98 | 0.99 | -2.08 | 0.16 | 0.56 | 0.49 |
| **Warthog** | *Phacochoerus africanus* 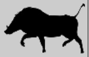 | H | NT | 0.79 | 0.79 | 0.99 | 1.00 | -1 | 0.15 | 0.56 | 0.52 |
| **Springbok** | *Antidorcas marsupialis* 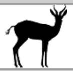 | H | NT | 0.76 | 0.78 | 0.99 | 0.99 | -7.93 | 0.19 | 0.51 | 0.77 |
| **Roan** | *Hippotragus equinus* 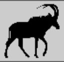 | H | NT | 0.69 | 0.68 | 0.80 | 0.99 | -10.09 | 0.13 | 0.54 | 0.82 |
| **Black wildebeest** | *Connochaetes gnou* 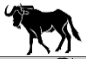 | H | NT | 0.74 | 0.76 | 0.96 | 0.99 | -5.37 | 0.23 | 0.47 | 0.86 |
| **Burchells zebra** | *Equus quagga* 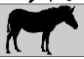 | H | NT | 0.76 | 0.78 | 0.99 | 1.00 | 4.25 | 0.13 | 0.53 | 0.92 |
| **Waterbuck** | *Kobusellipsiprymnus* 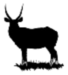 | H | NT | 0.81 | 0.72 | 0.98 | 0.99 | -26.08 | 0.15 | 0.51 | 0.97 |
| **Blue wildebeest** | *Connochaetes taurinus* 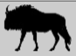 | H | NT | 0.77 | 0.80 | 1.00 | 0.97 | 6.19 | 0.09 | 0.54 | 1.06 |
| **Giraffe** | *Giraffa camelopardalis* 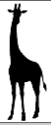 | H | T | 0.80 | 0.82 | 0.99 | 0.99 | 5.78 | 0.10 | 0.54 | 1.07 |
| **Mountain reedbuck** | *Redunca fulvorufula* 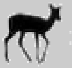 | H | T | 0.71 | 0.70 | 0.99 | 1.00 | -3.94 | 0.13 | 0.50 | 1.21 |
| **Sable antelope** | *Hippotragus niger* 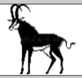 | H | NT | 0.73 | 0.76 | 0.99 | 0.97 | -1.94 | 0.13 | 0.50 | 1.26 |
| **Kudu** | *Tragelaphus strepsiceros* 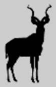 | H | NT | 0.79 | 0.81 | 1.00 | 1.00 | 5.02 | 0.09 | 0.52 | 1.3 |
| **Bontebok** | *Damaliscus pygargus phillipsi* 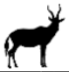 | H | T | 0.69 | 0.63 | 0.93 | 0.88 | -4.78 | 0.25 | 0.41 | 1.42 |
| **Impala** | *Aepyceros melampus* 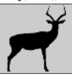 | H | NT | 0.80 | 0.72 | 0.99 | 1.00 | -12.79 | 0.11 | 0.47 | 1.7 |
