## Supplementary Information 2 - HSMs results and maps for "Shifted distribution baselines: neglecting long-term biodiversity records risks overlooking potentially suitable habitat for conservation management"

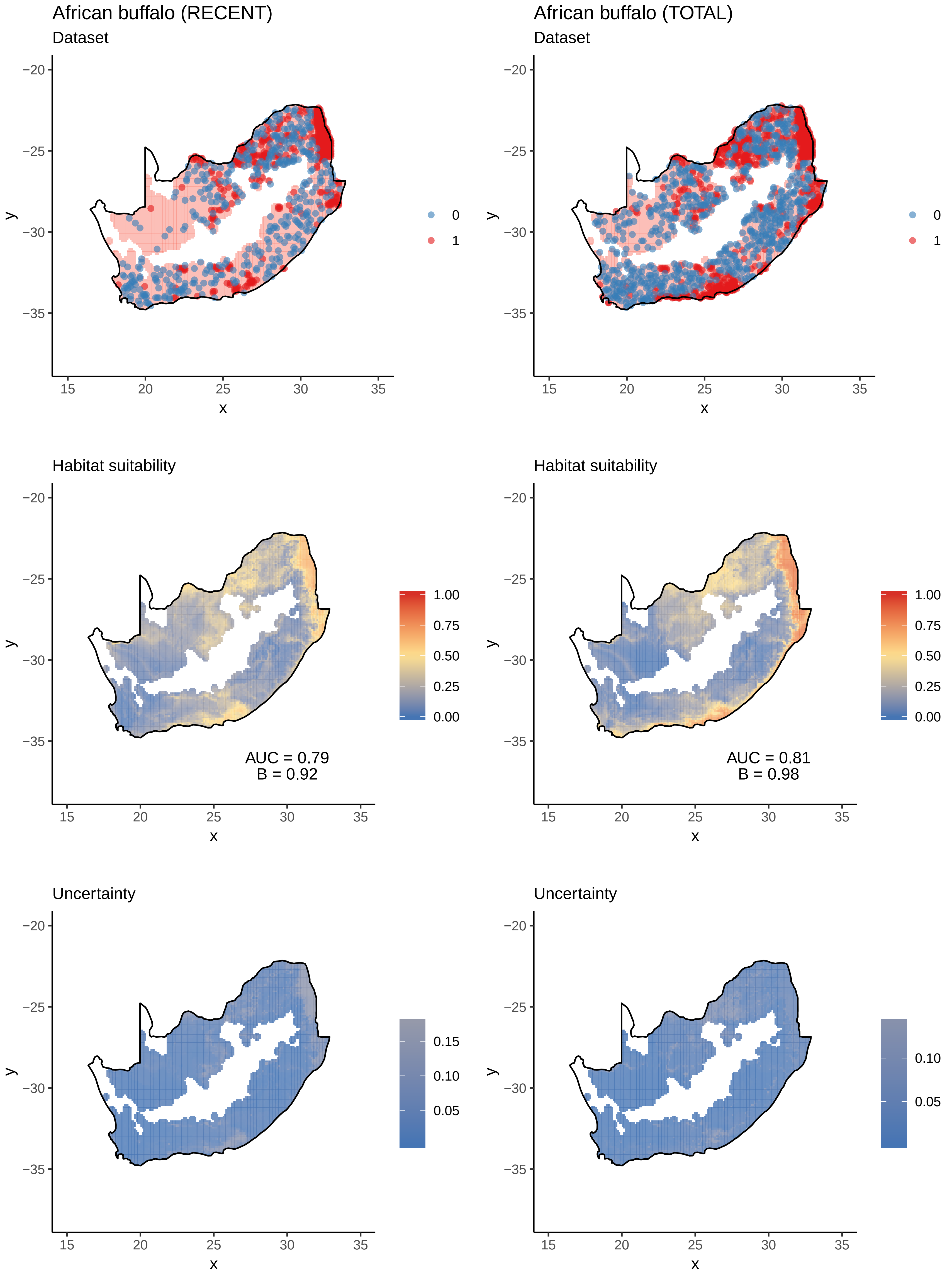

African buffalo (RECENT)
Dataset
African buffalo (TOTAL)
Dataset
0
1
Habitat suitability
AUC = 0.79
B = 0.92
AUC = 0.81
B = 0.98
1.00
0.75
0.50
0.25
0.00
Uncertainty
0.15
0.10
0.05
x
y

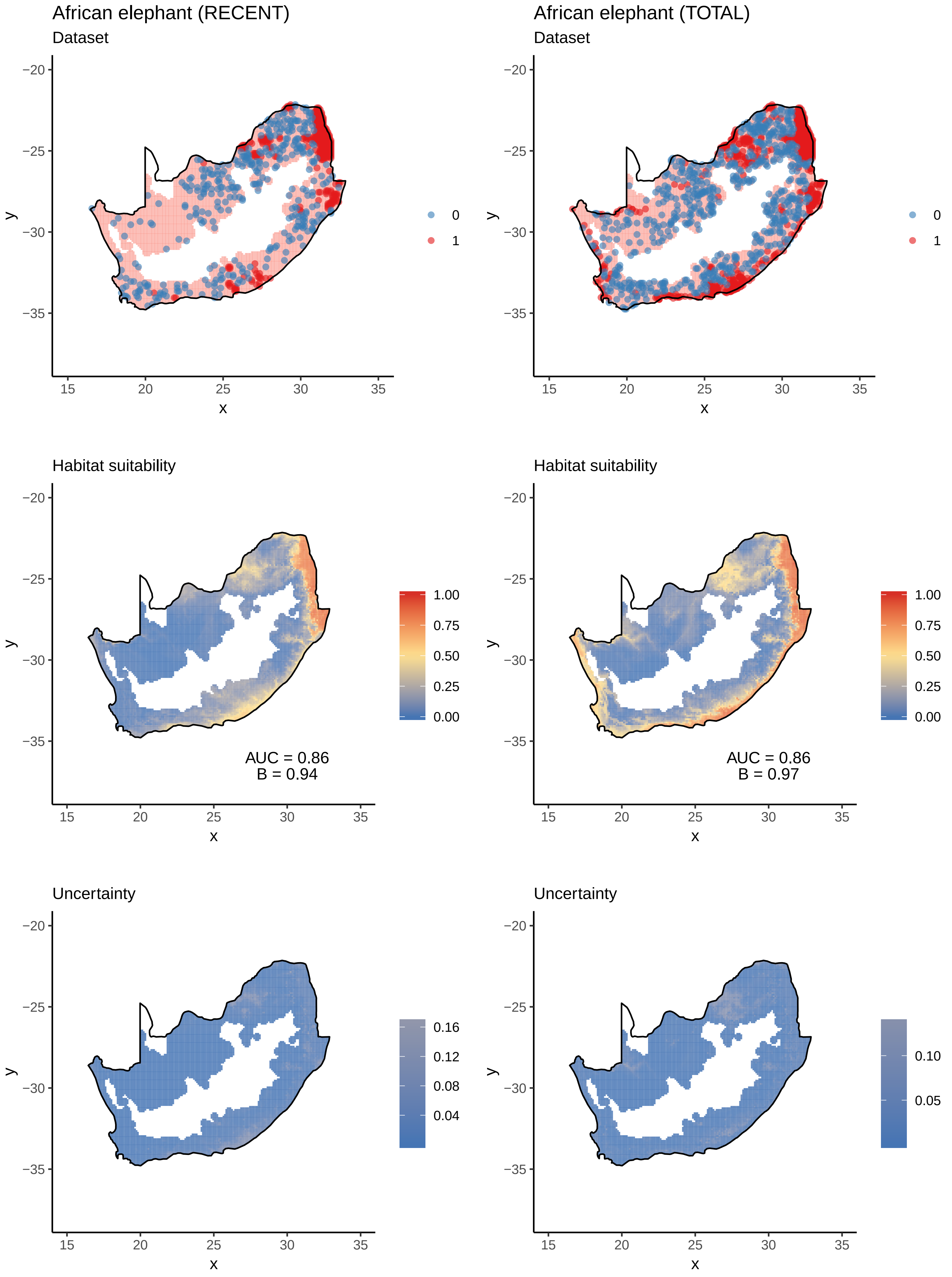

African elephant (RECENT)
Dataset
African elephant (TOTAL)
Dataset
0
1
Habitat suitability
Habitat suitability
1.00
0.75
0.50
0.25
0.00
AUC = 0.86
B = 0.94
AUC = 0.86
B = 0.97
Uncertainty
Uncertainty
0.16
0.12
0.08
0.04
0.10
0.05
−20
−25
−30
−35
15
20
25
30
35
x
y

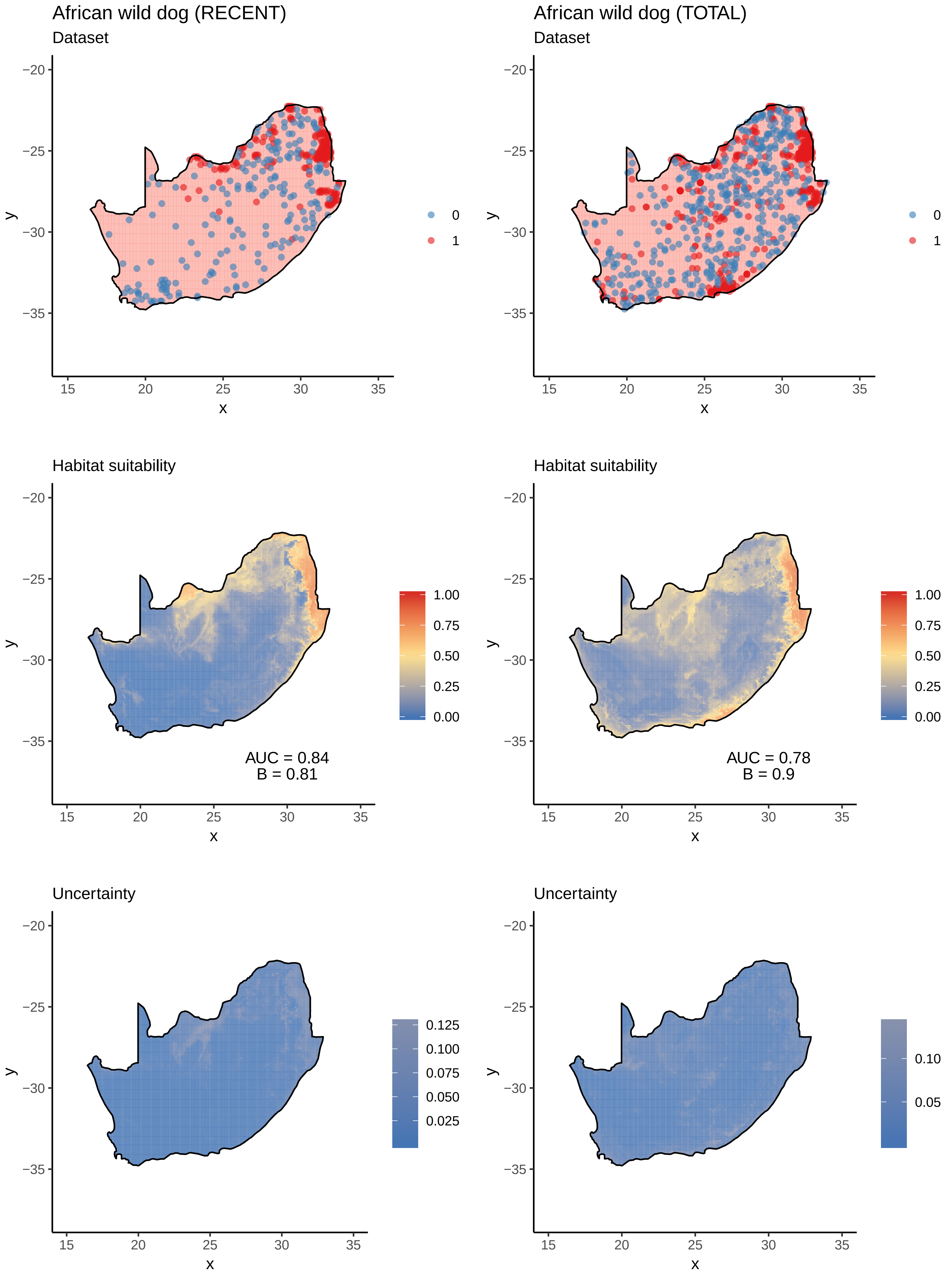
African wild dog (RECENT)
Dataset
African wild dog (TOTAL)
Dataset
Habitat suitability
AUC = 0.84
B = 0.81
Habitat suitability
AUC = 0.78
B = 0.9
Uncertainty
Uncertainty
x
y
0
1
1.00
0.75
0.50
0.25
0.00
0.125
0.100
0.075
0.050
0.025
0.10
0.05

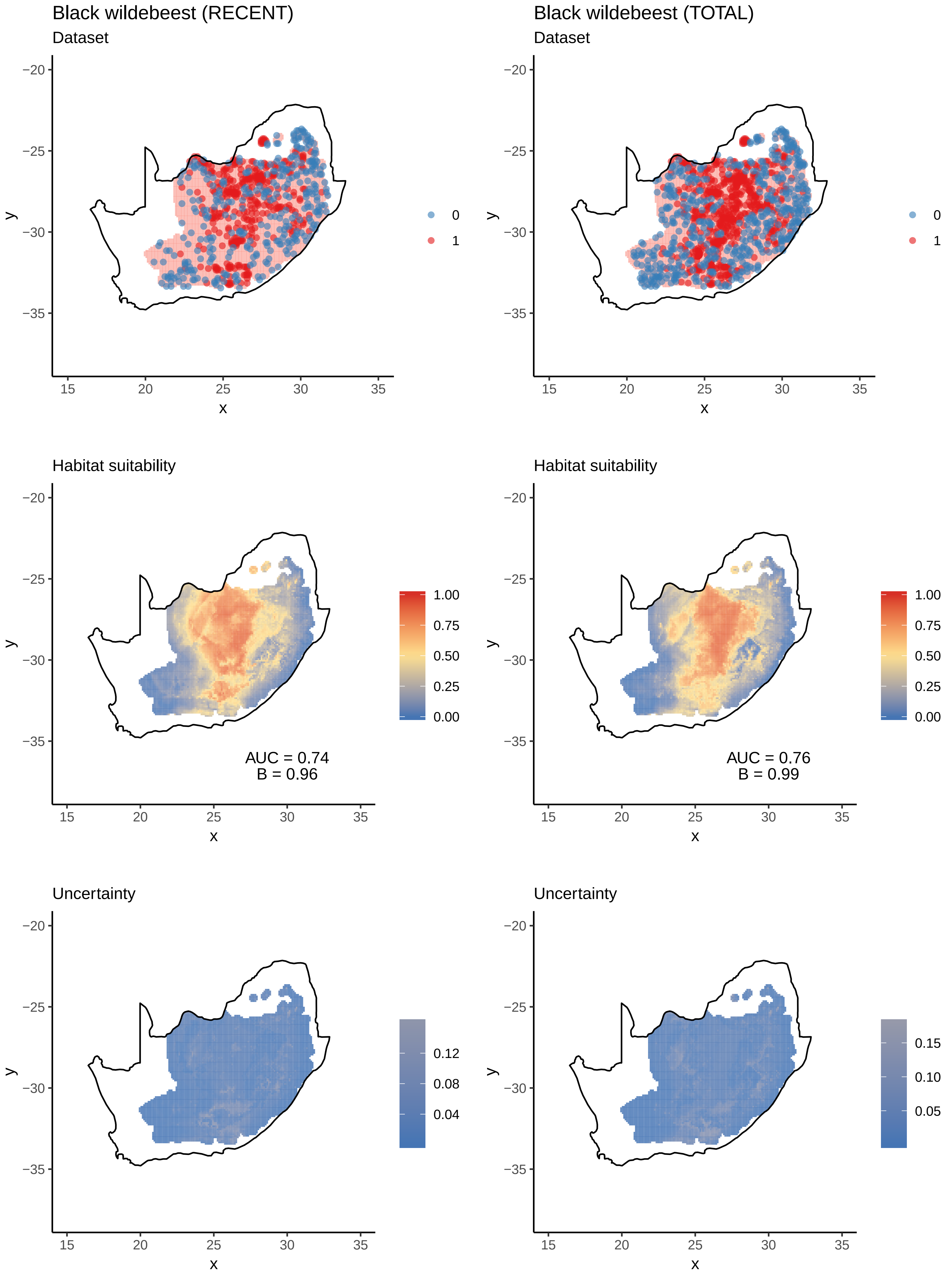

Black wildebeest (RECENT)
Dataset
Black wildebeest (TOTAL)
Dataset
Habitat suitability
AUC = 0.74
B = 0.96
Habitat suitability
AUC = 0.76
B = 0.99
Uncertainty
Uncertainty

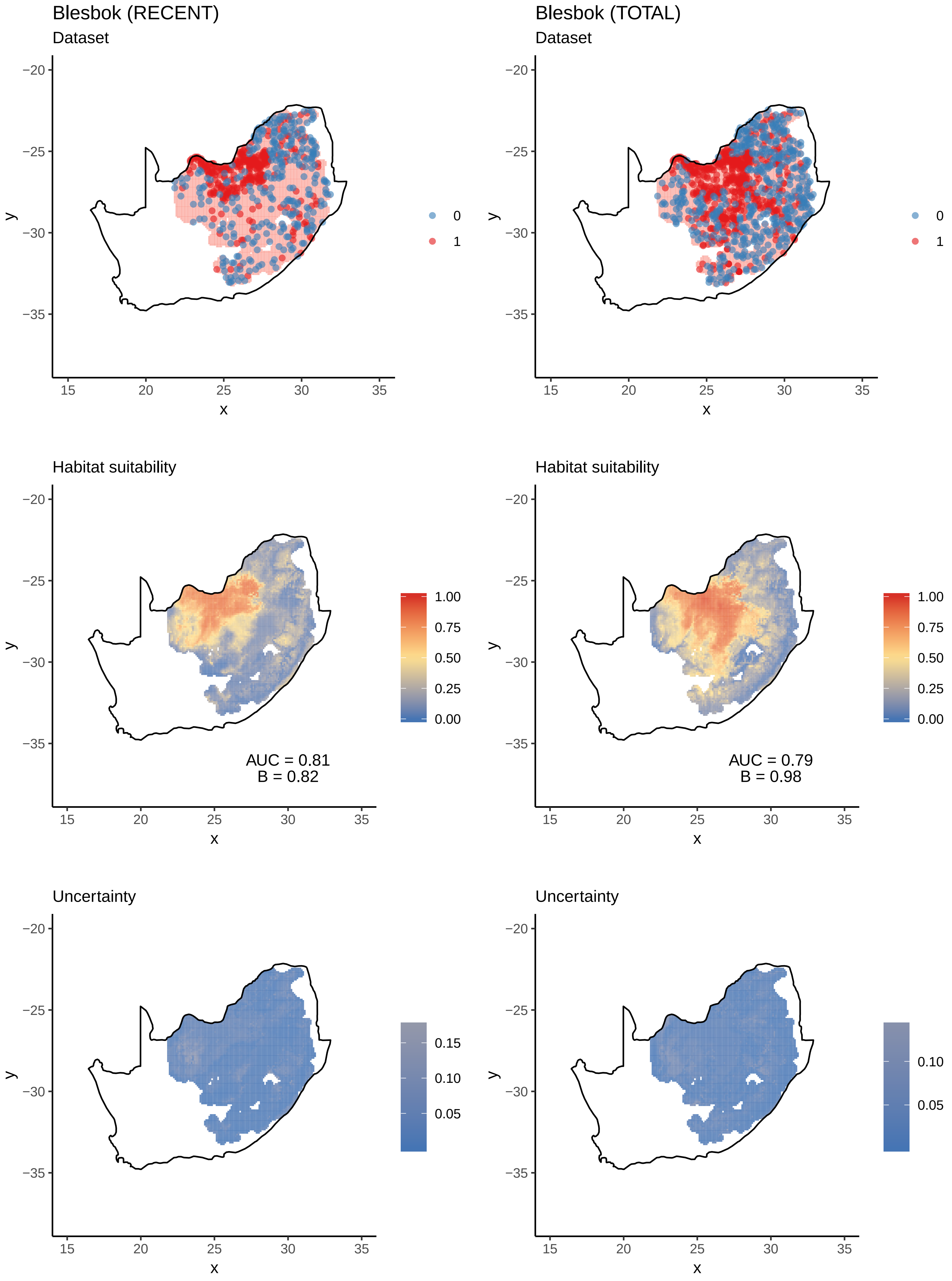

Blesbok (RECENT)
Dataset
Blesbok (TOTAL)
Dataset
0
1
Habitat suitability
AUC = 0.81
B = 0.82
AUC = 0.79
B = 0.98
1.00
0.75
0.50
0.25
0.00
Uncertainty
0.15
0.10
0.05
x
y

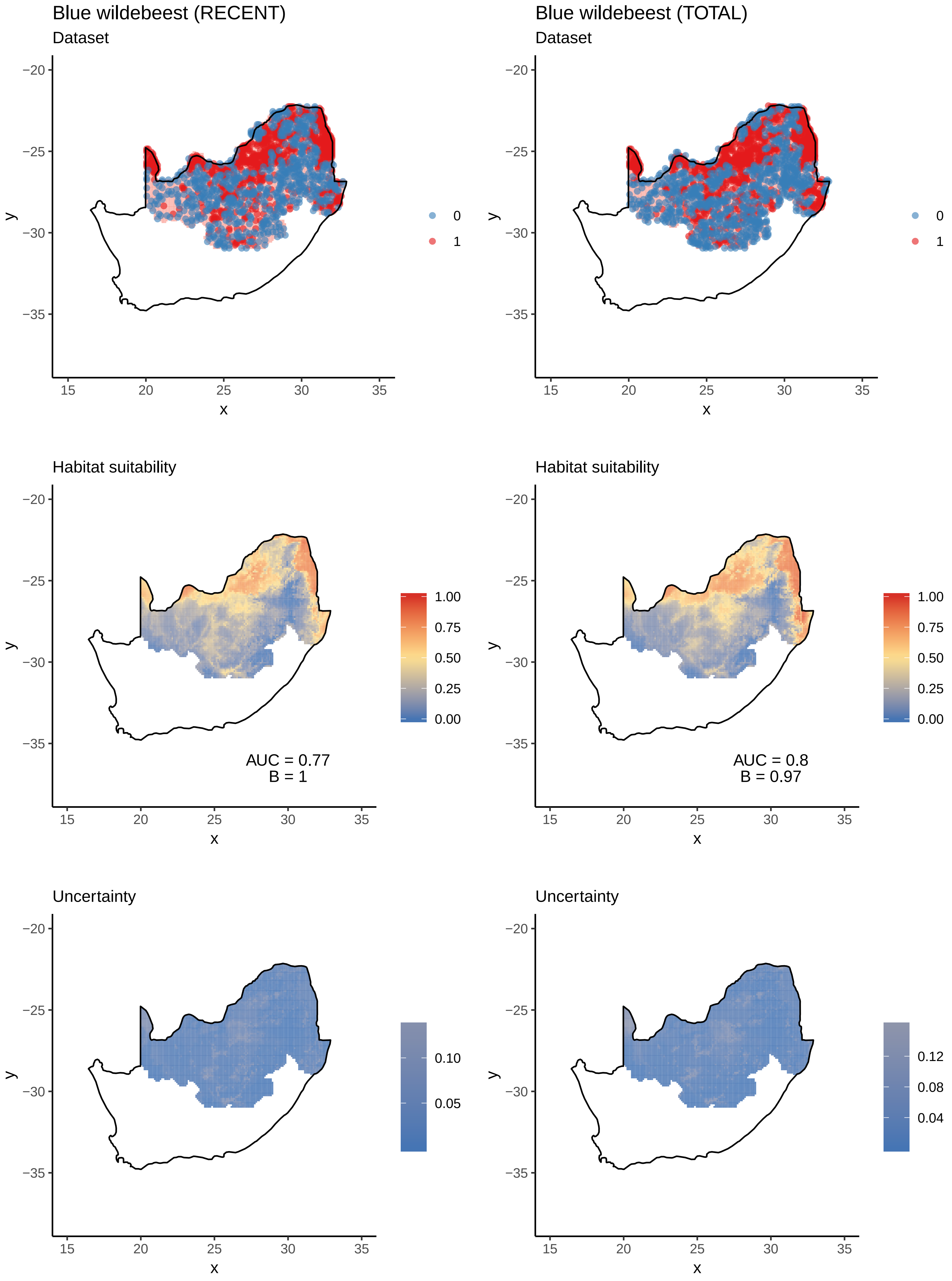

Blue wildebeest (RECENT)
Dataset
Blue wildebeest (TOTAL)
Dataset
0
1
Habitat suitability
AUC = 0.77
B = 1
AUC = 0.8
B = 0.97
1.00
0.75
0.50
0.25
0.00
Uncertainty
0.10
0.05
0.12
0.08
0.04
−20
−25
−30
−35
15
20
25
30
35
x
y

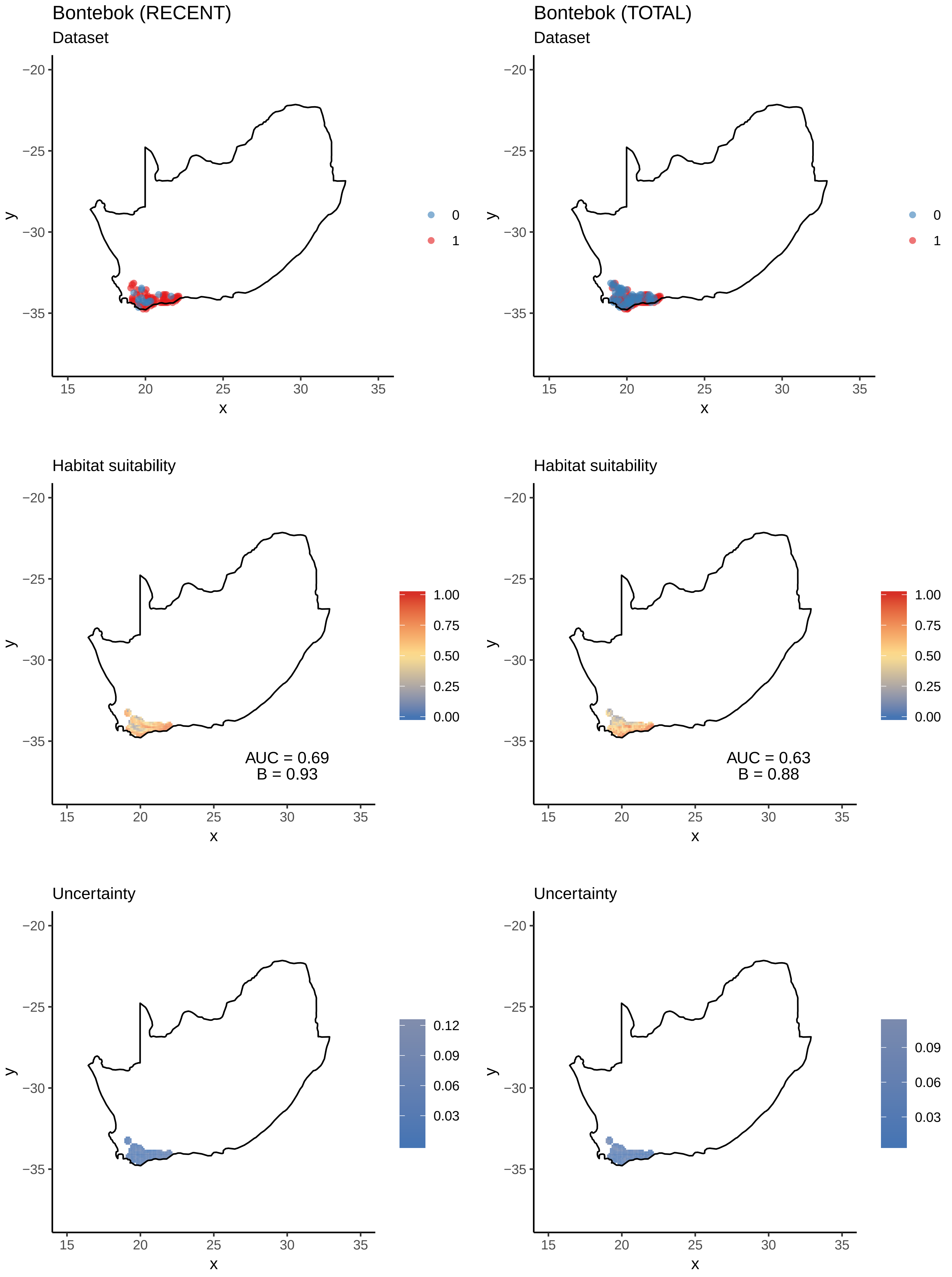
Bontebok (RECENT)
Dataset
Bontebok (TOTAL)
Dataset
Habitat suitability
AUC = 0.69
B = 0.93
Habitat suitability
AUC = 0.63
B = 0.88
Uncertainty
Uncertainty
x
y
0
1
1.00
0.75
0.50
0.25
0.00
0.12
0.09
0.06
0.03

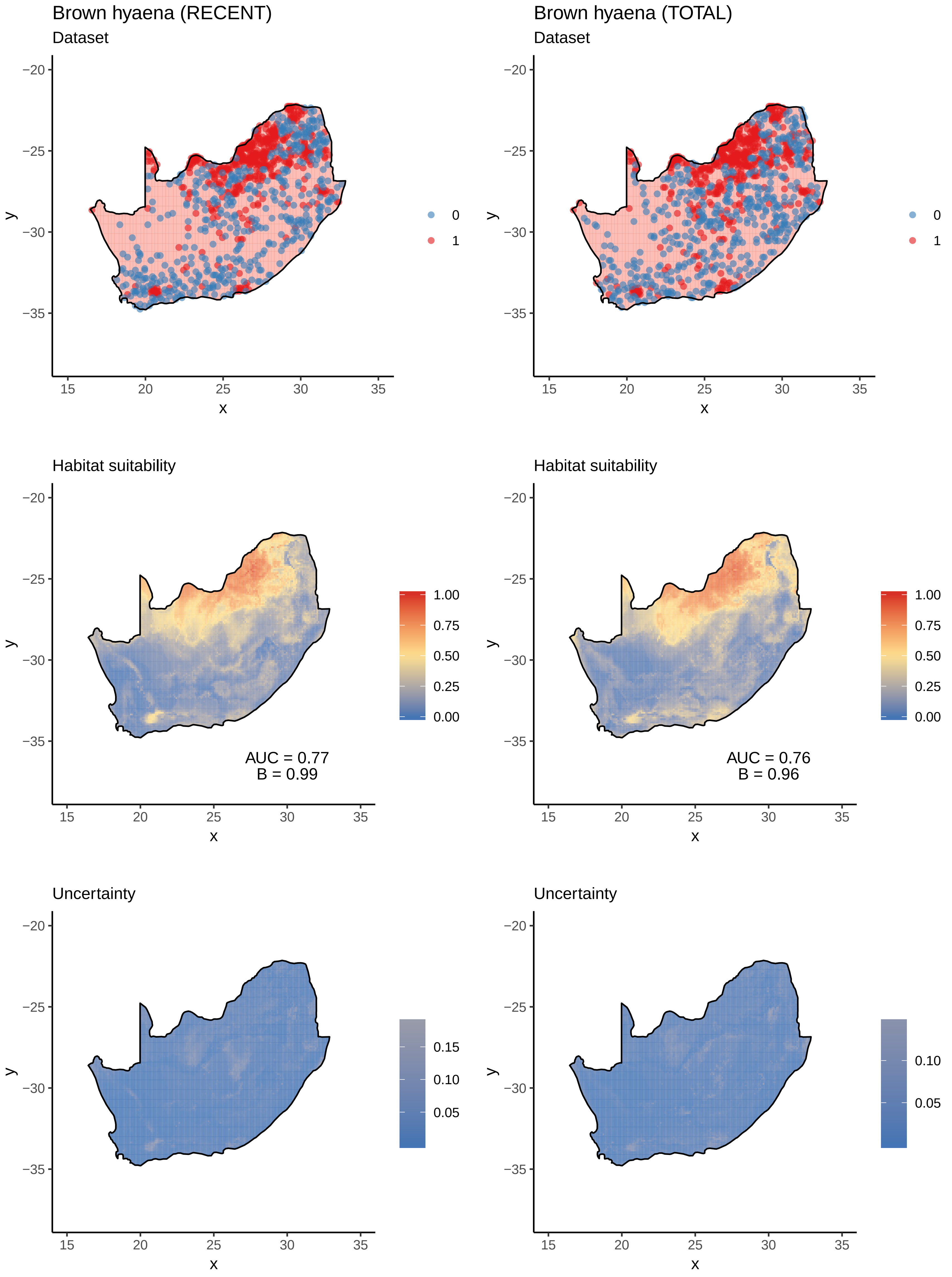

Brown hyaena (RECENT)
Dataset
Brown hyaena (TOTAL)
Dataset
0
1
Habitat suitability
AUC = 0.77
B = 0.99
AUC = 0.76
B = 0.96
1.00
0.75
0.50
0.25
0.00
Uncertainty
0.15
0.10
0.05
x
y
−20
−25
−30
−35
15
20
25
30
35

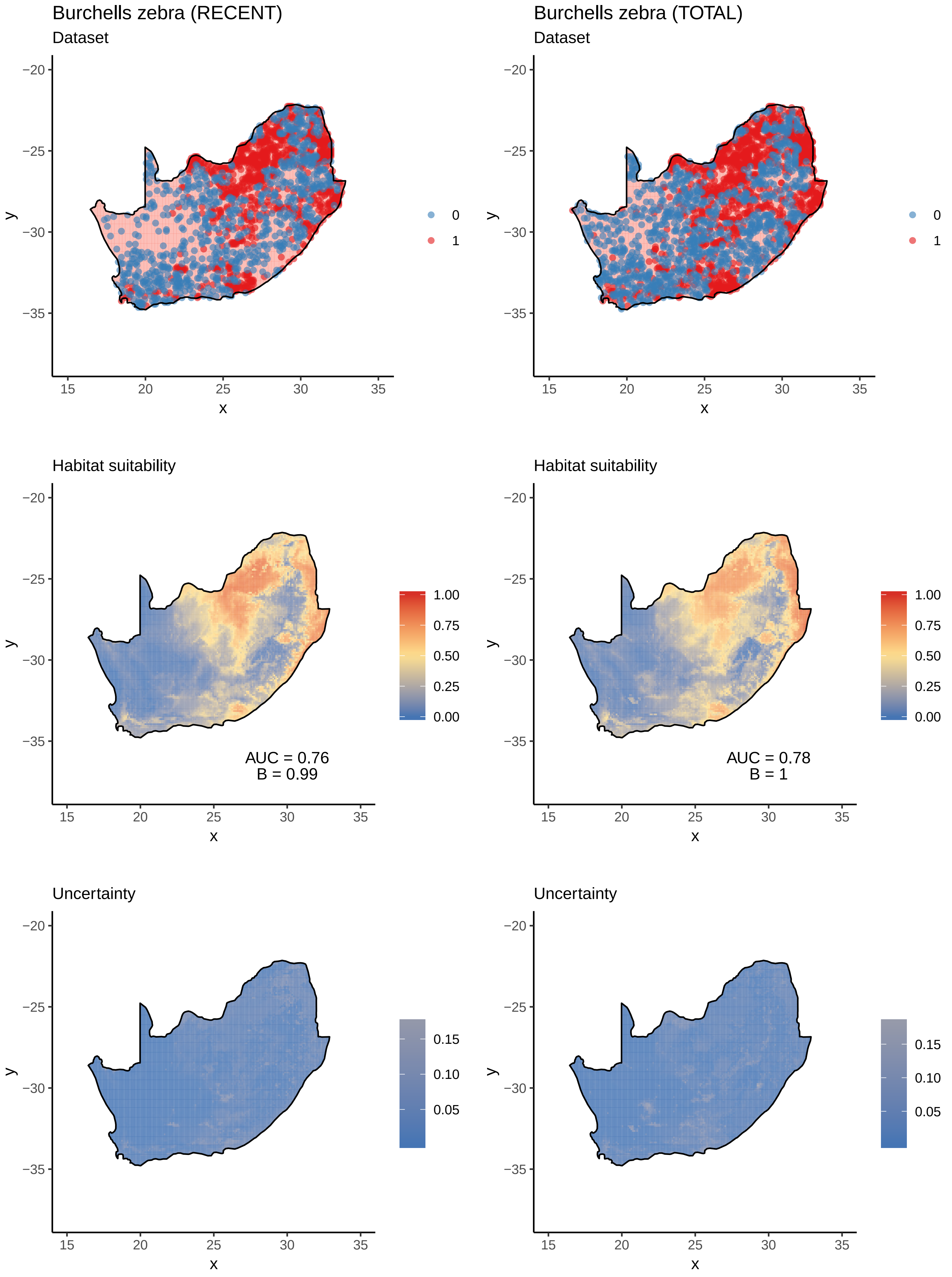

Burchells zebra (RECENT)
Dataset
Burchells zebra (TOTAL)
Dataset
Habitat suitability
AUC = 0.76
B = 0.99
Habitat suitability
AUC = 0.78
B = 1
Uncertainty
Uncertainty
x
y
0
1
1.00
0.75
0.50
0.25
0.00
0.15
0.10
0.05

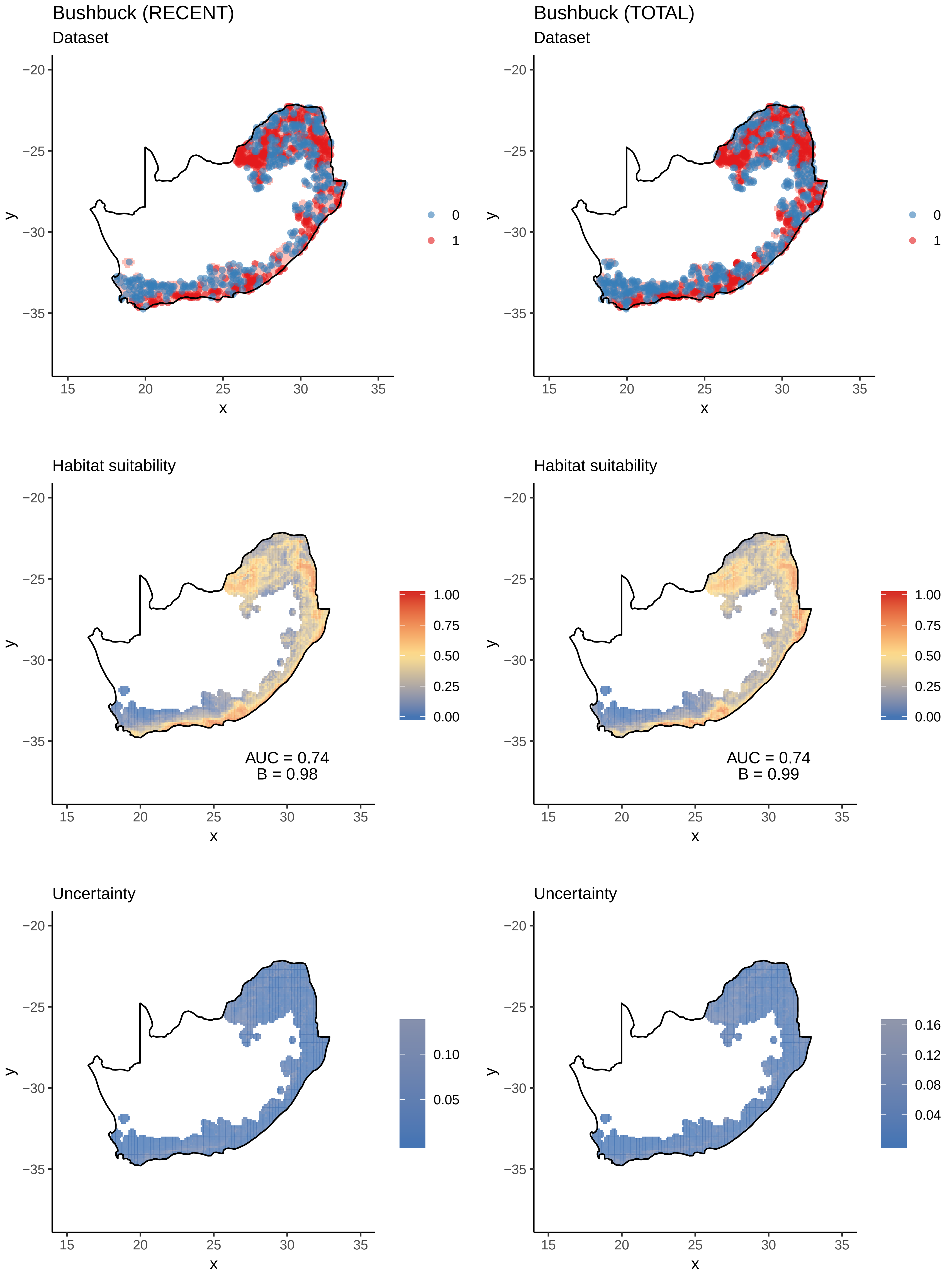

Bushbuck (RECENT)
Dataset
Bushbuck (TOTAL)
Dataset
0
1
Habitat suitability
AUC = 0.74
B = 0.98
Habitat suitability
AUC = 0.74
B = 0.99
1.00
0.75
0.50
0.25
0.00
Uncertainty
0.10
0.05
Uncertainty
0.16
0.12
0.08
0.04
x
y
15
20
25
30
35
−20
−25
−30
−35

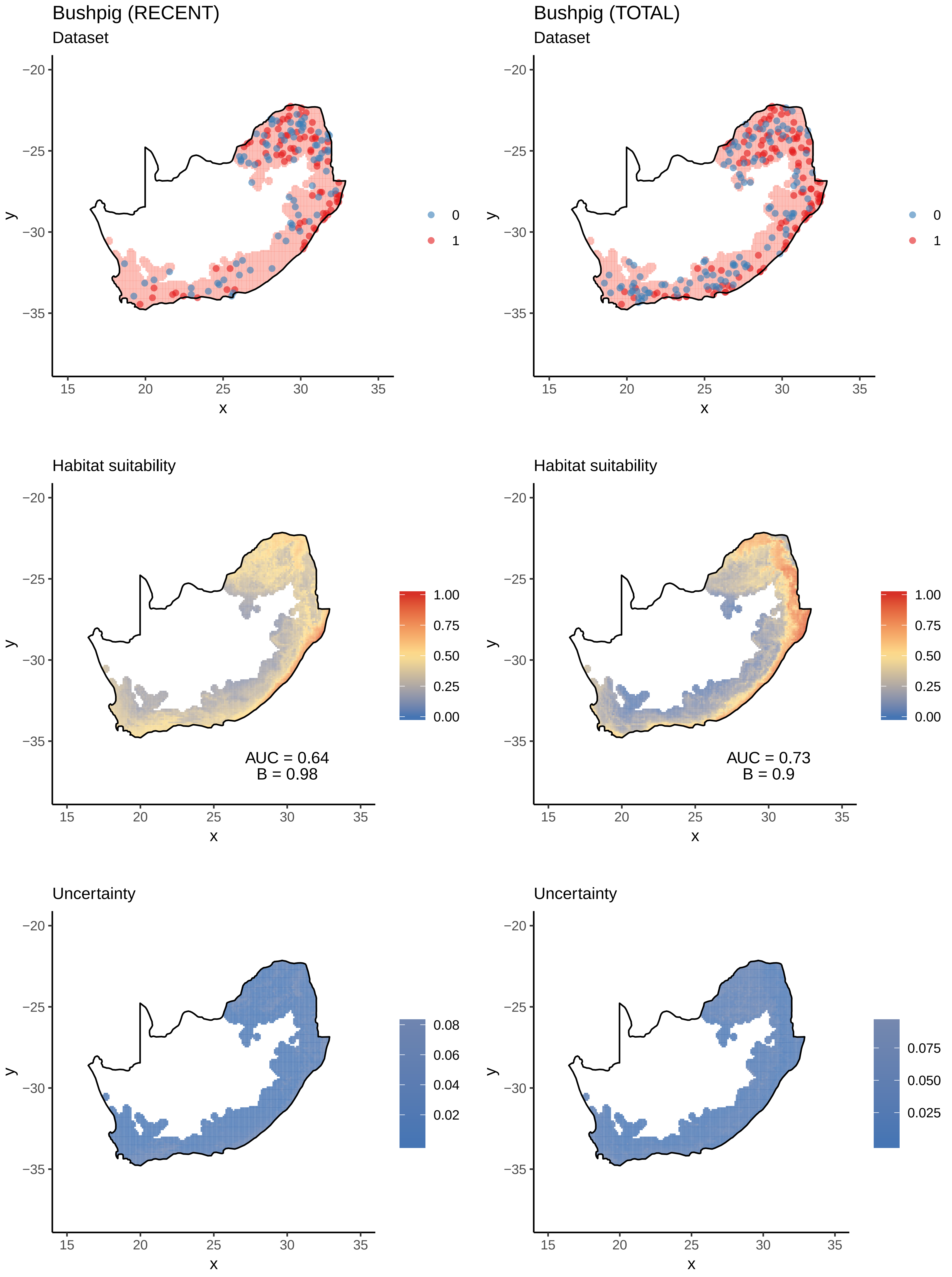
Bushpig (RECENT)
Dataset
Bushpig (TOTAL)
Dataset
Habitat suitability
AUC = 0.64
B = 0.98
Habitat suitability
AUC = 0.73
B = 0.9
Uncertainty
Uncertainty
x
y

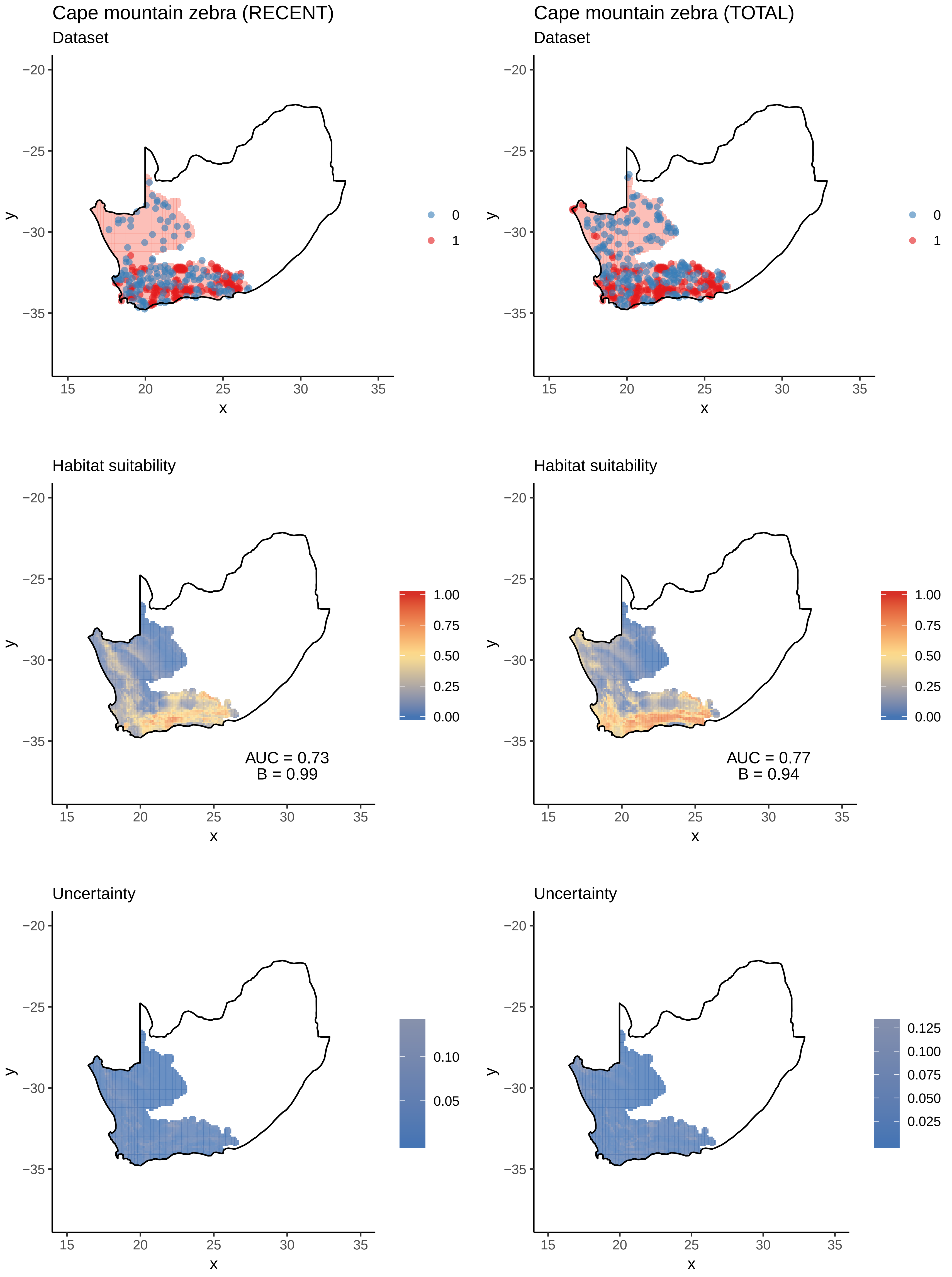
Cape mountain zebra (RECENT)
Dataset
Cape mountain zebra (TOTAL)
Dataset
Habitat suitability
AUC = 0.73
B = 0.99
Habitat suitability
AUC = 0.77
B = 0.94
Uncertainty
Uncertainty
x
y
0
1

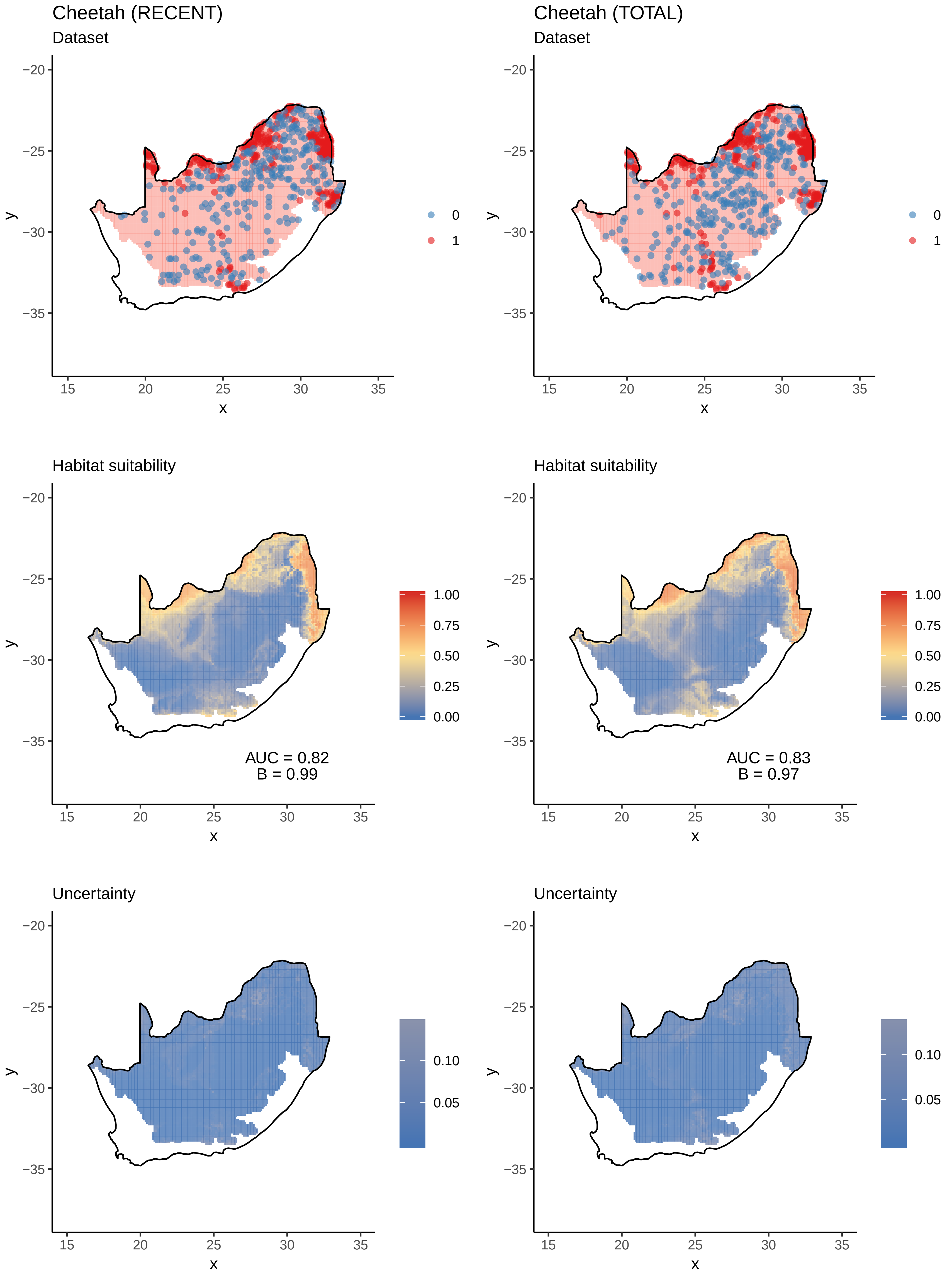
Cheetah (RECENT)
Dataset
Cheetah (TOTAL)
Dataset
0
1
Habitat suitability
AUC = 0.82
B = 0.99
AUC = 0.83
B = 0.97
1.00
0.75
0.50
0.25
0.00
Uncertainty
0.10
0.05
x
y

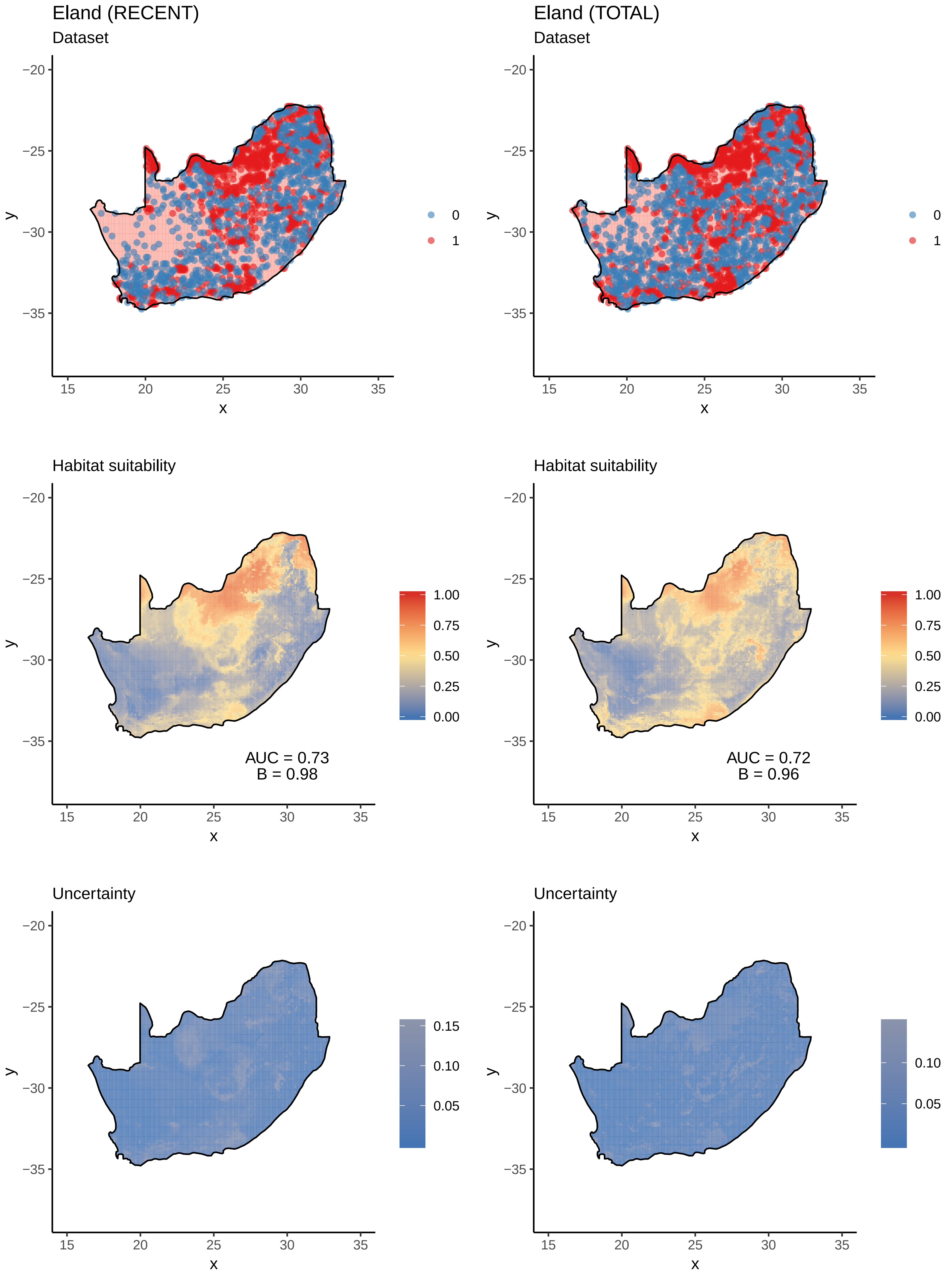

Eland (RECENT)
Dataset
Eland (TOTAL)
Dataset
0
1
Habitat suitability
AUC = 0.73
B = 0.98
AUC = 0.72
B = 0.96
1.00
0.75
0.50
0.25
0.00
Uncertainty
0.15
0.10
0.05
−20
−25
−30
−35
15
20
25
30
35
x
y

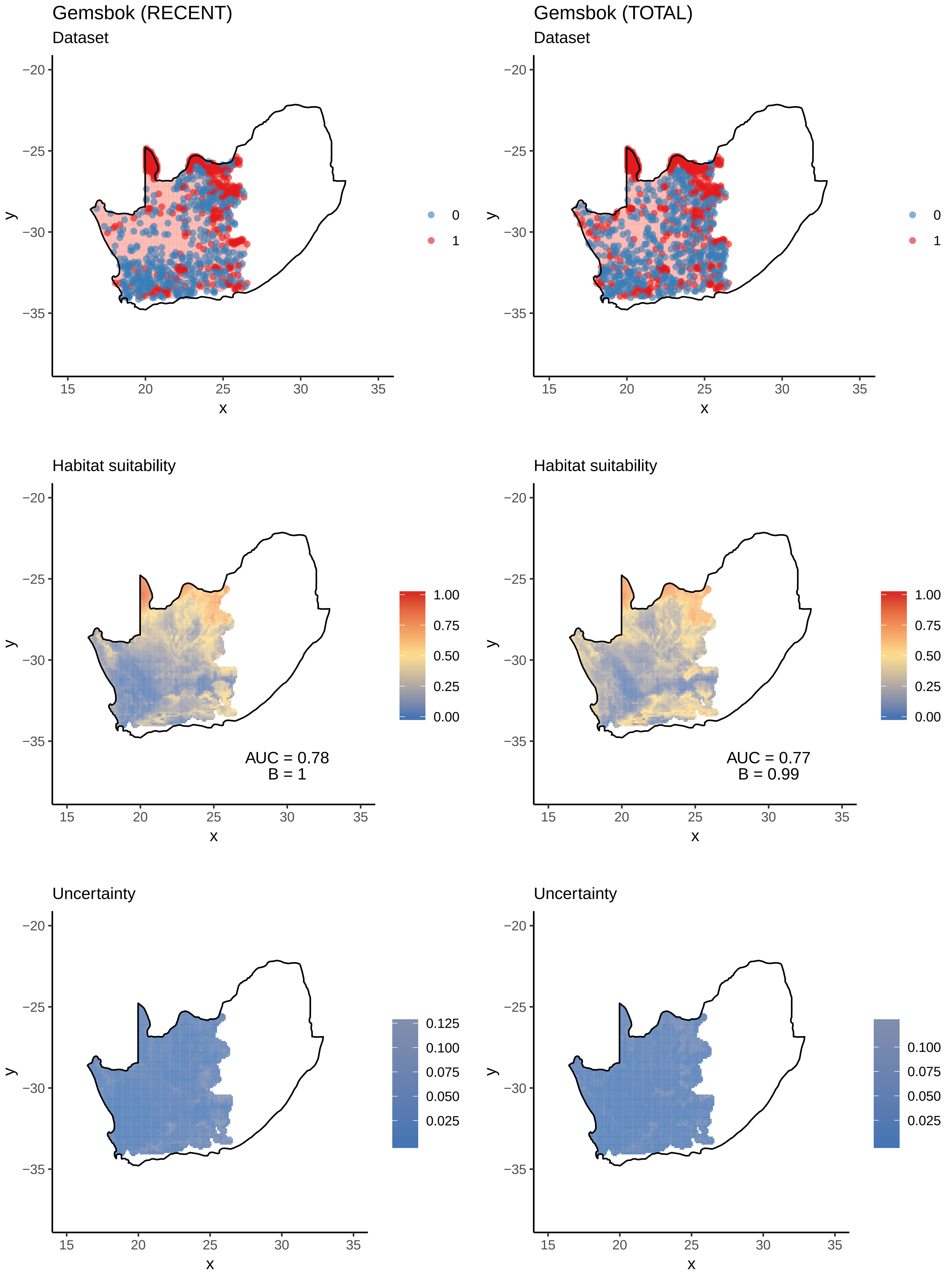
Gemsbok (RECENT)
Dataset
Gemsbok (TOTAL)
Dataset
Habitat suitability
AUC = 0.78
B = 1
Habitat suitability
AUC = 0.77
B = 0.99
Uncertainty
Uncertainty
x
y
0
1
1.00
0.75
0.50
0.25
0.00
0.125
0.100
0.075
0.050
0.025
15
20
25
30
35
−20
−25
−30
−35

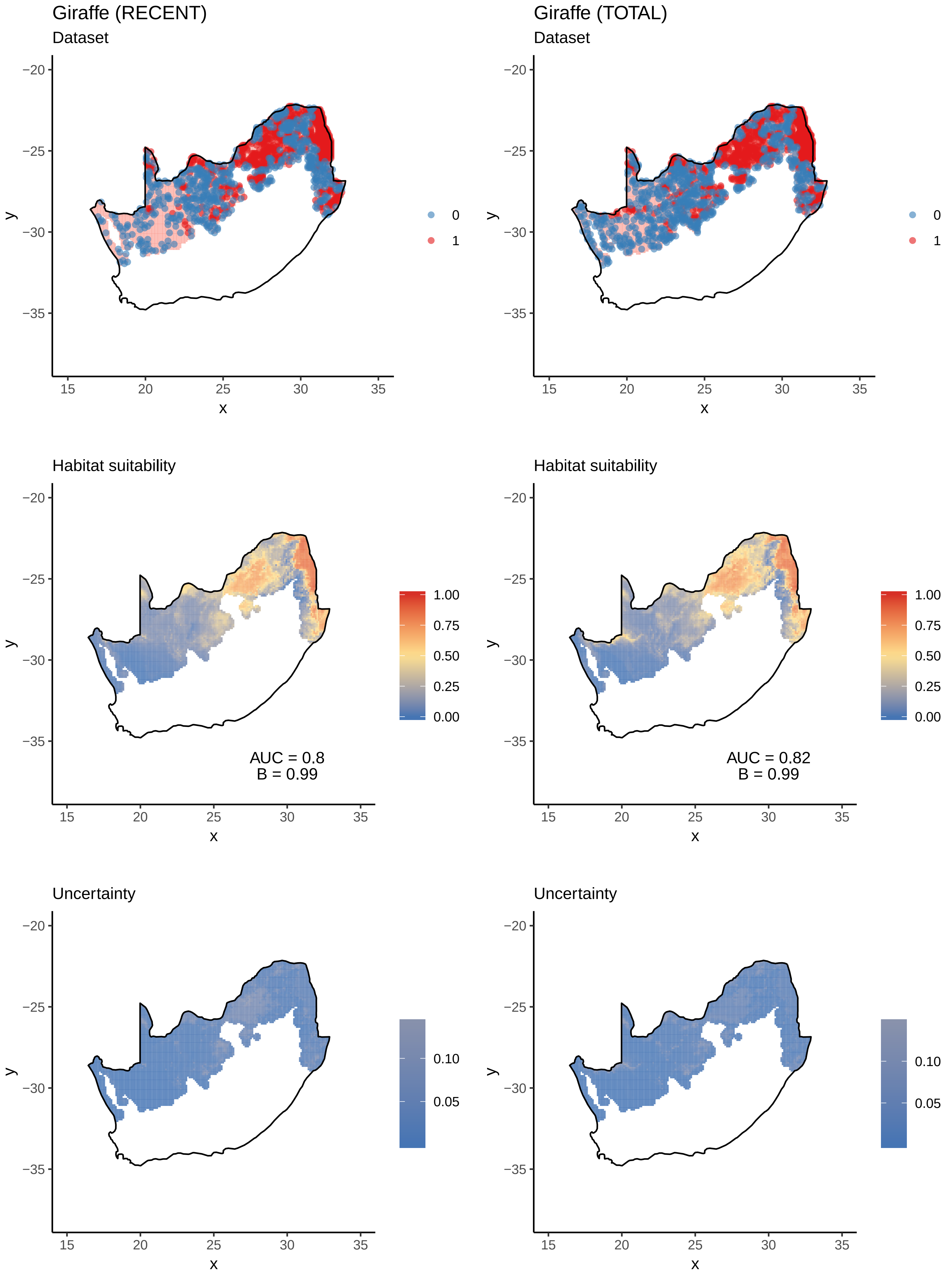

Giraffe (RECENT)
Dataset
Giraffe (TOTAL)
Dataset
0
1
Habitat suitability
AUC = 0.8
B = 0.99
AUC = 0.82
B = 0.99
1.00
0.75
0.50
0.25
0.00
Uncertainty
0.10
0.05
x
y

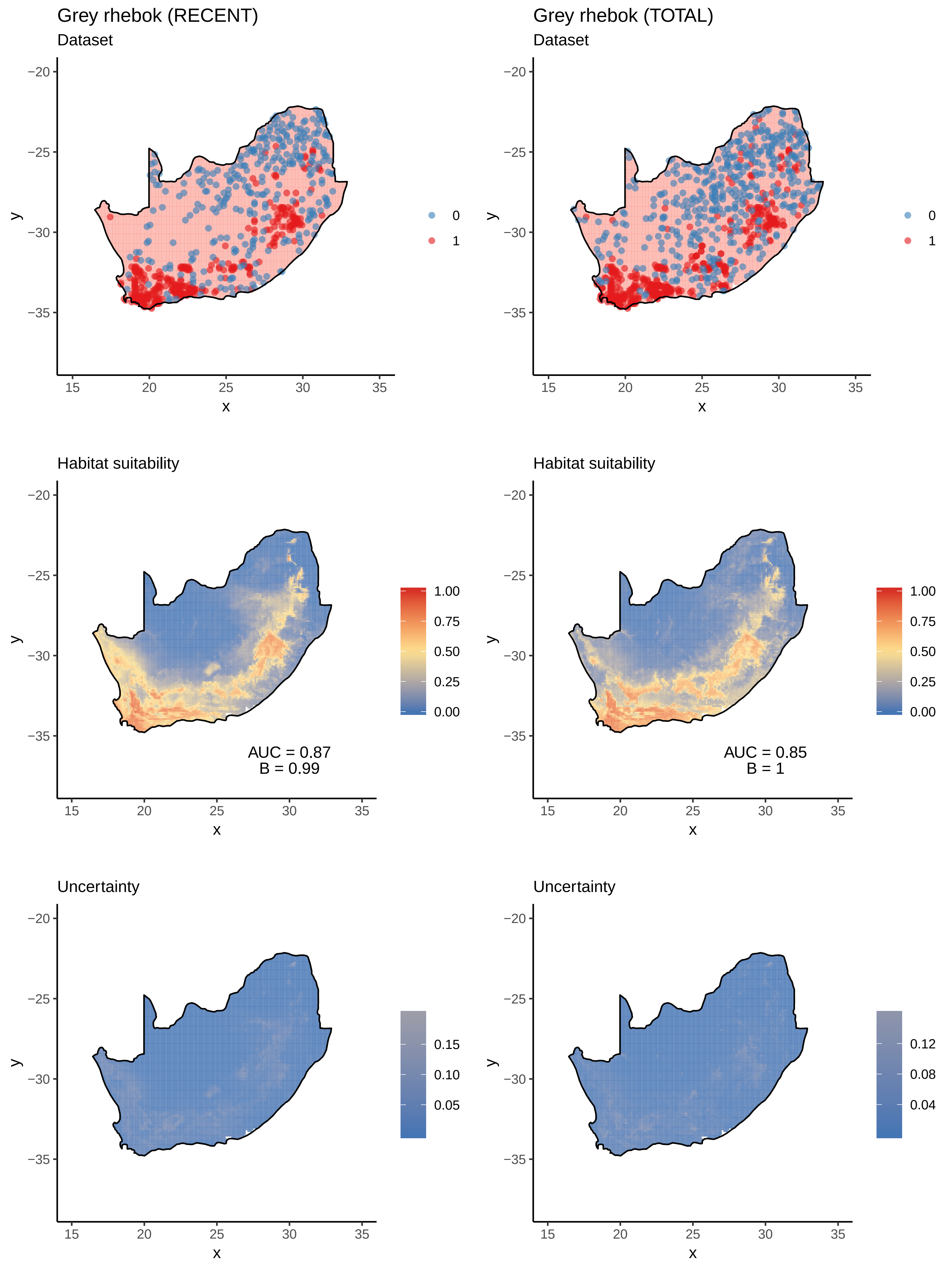

Grey rhebok (RECENT)
Dataset
Grey rhebok (TOTAL)
Dataset
Habitat suitability
AUC = 0.87
B = 0.99
Habitat suitability
AUC = 0.85
B = 1
Uncertainty
Uncertainty
x
y
0
1
1.00
0.75
0.50
0.25
0.00
0.15
0.10
0.05
0.12
0.08
0.04

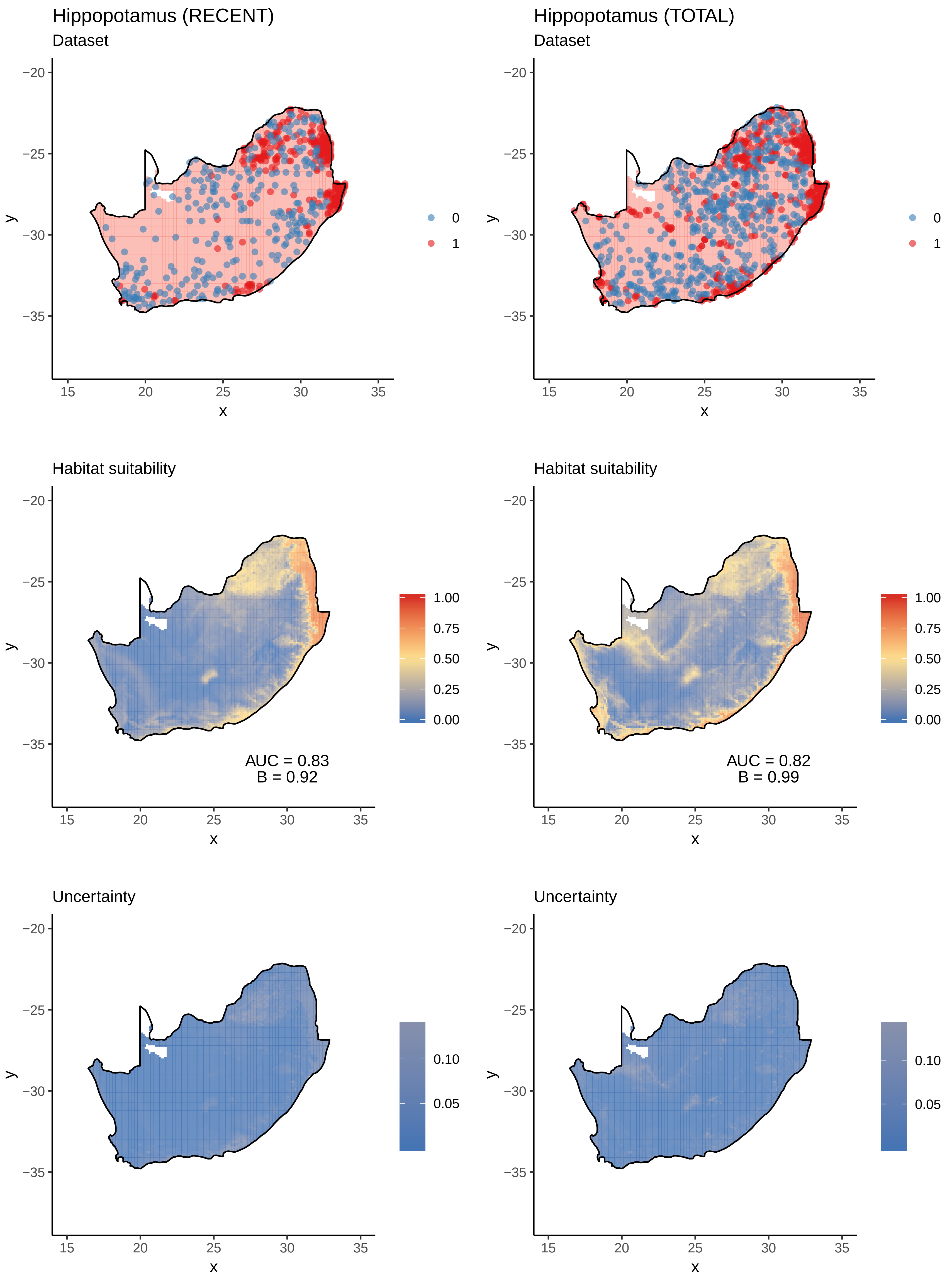
Hippopotamus (RECENT)
Dataset
Hippopotamus (TOTAL)
Dataset
0
1
Habitat suitability
AUC = 0.83
B = 0.92
AUC = 0.82
B = 0.99
1.00
0.75
0.50
0.25
0.00
Uncertainty
0.10
0.05
x
y
15
20
25
30
35
−20
−25
−30
−35

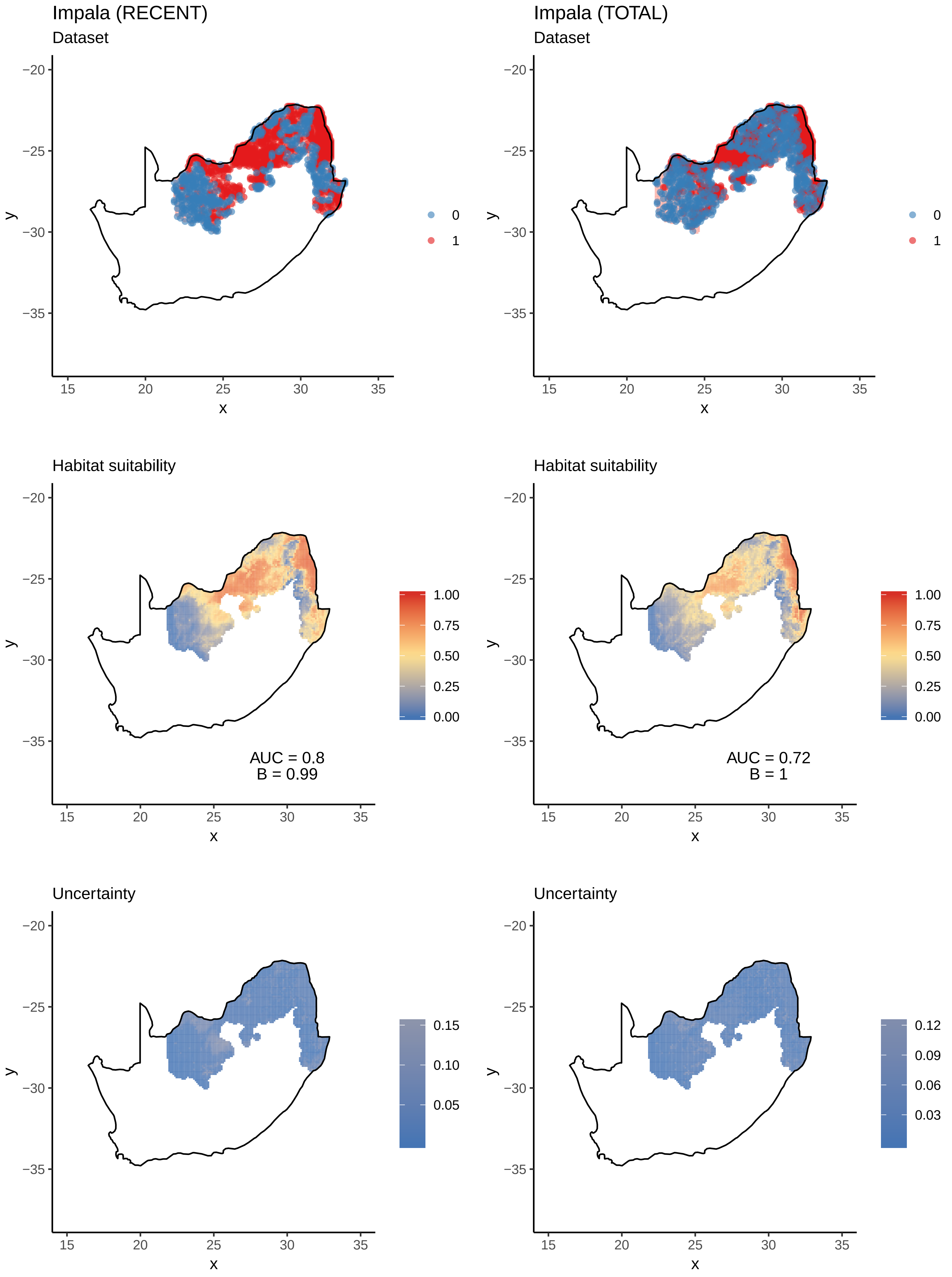
Impala (RECENT)
Dataset
Impala (TOTAL)
Dataset
0
1
Habitat suitability
AUC = 0.8
B = 0.99
AUC = 0.72
B = 1
1.00
0.75
0.50
0.25
0.00
Uncertainty
0.15
0.10
0.05
0.12
0.09
0.06
0.03
−20
−25
−30
−35
15
20
25
30
35
x
y

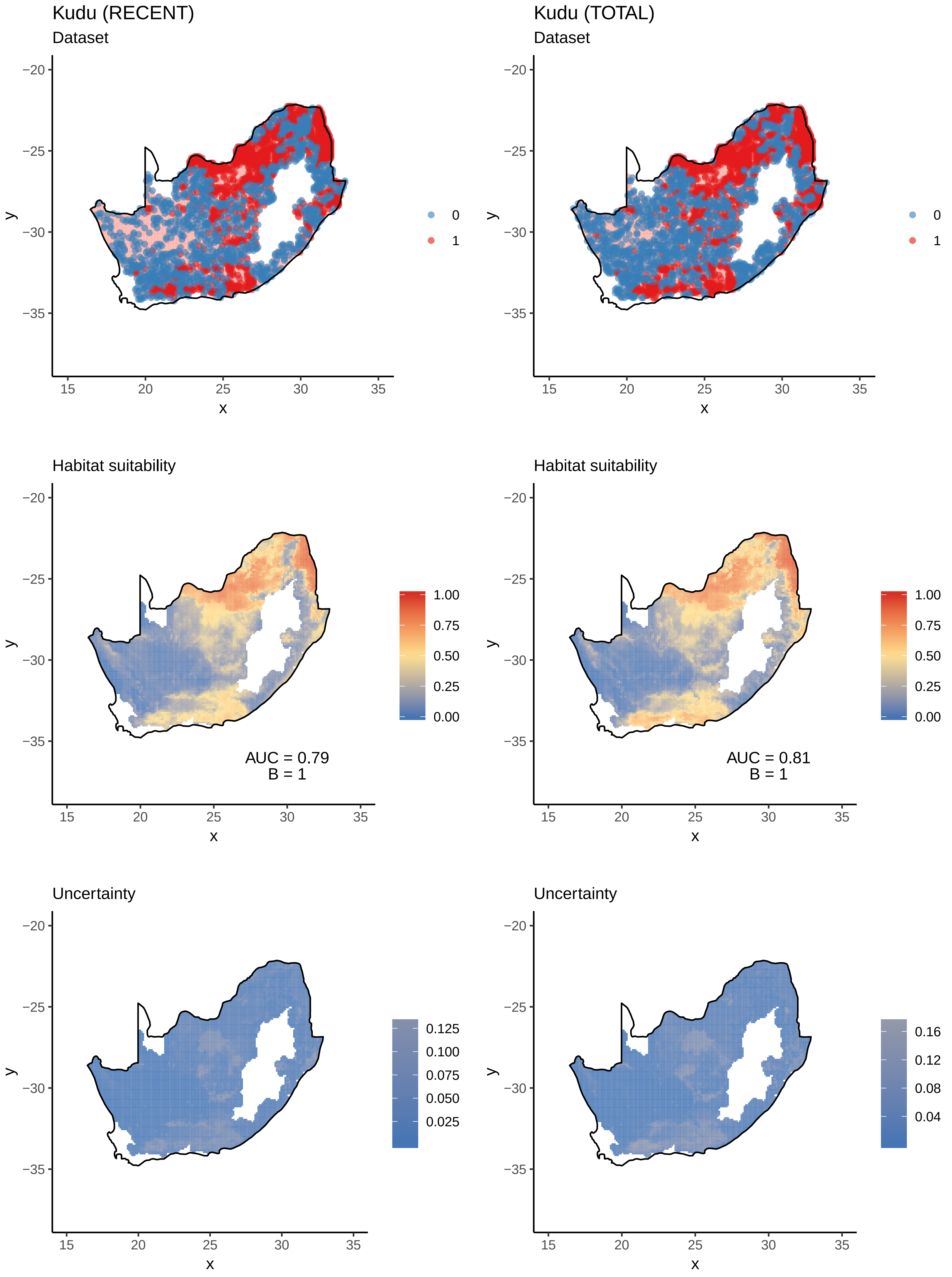
Kudu (RECENT)
Dataset
Kudu (TOTAL)
Dataset
Habitat suitability
AUC = 0.79
B = 1
AUC = 0.81
B = 1
Uncertainty
x
y
0
1
1.00
0.75
0.50
0.25
0.00
0.125
0.100
0.075
0.050
0.025
0.16
0.12
0.08
0.04

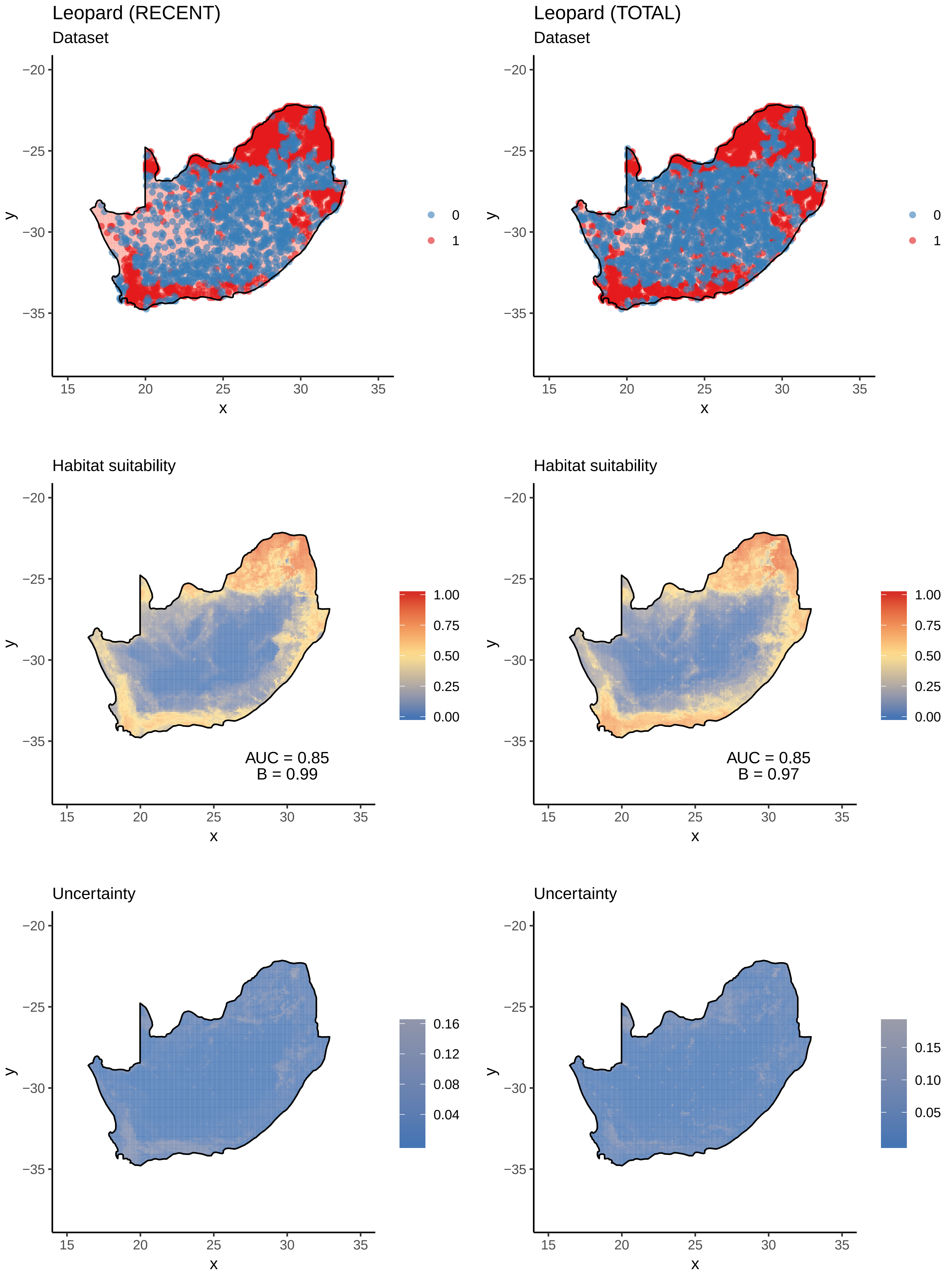

Leopard (RECENT)
Dataset
Leopard (TOTAL)
Dataset
Habitat suitability
AUC = 0.85
B = 0.99
Habitat suitability
AUC = 0.85
B = 0.97
Uncertainty
Uncertainty

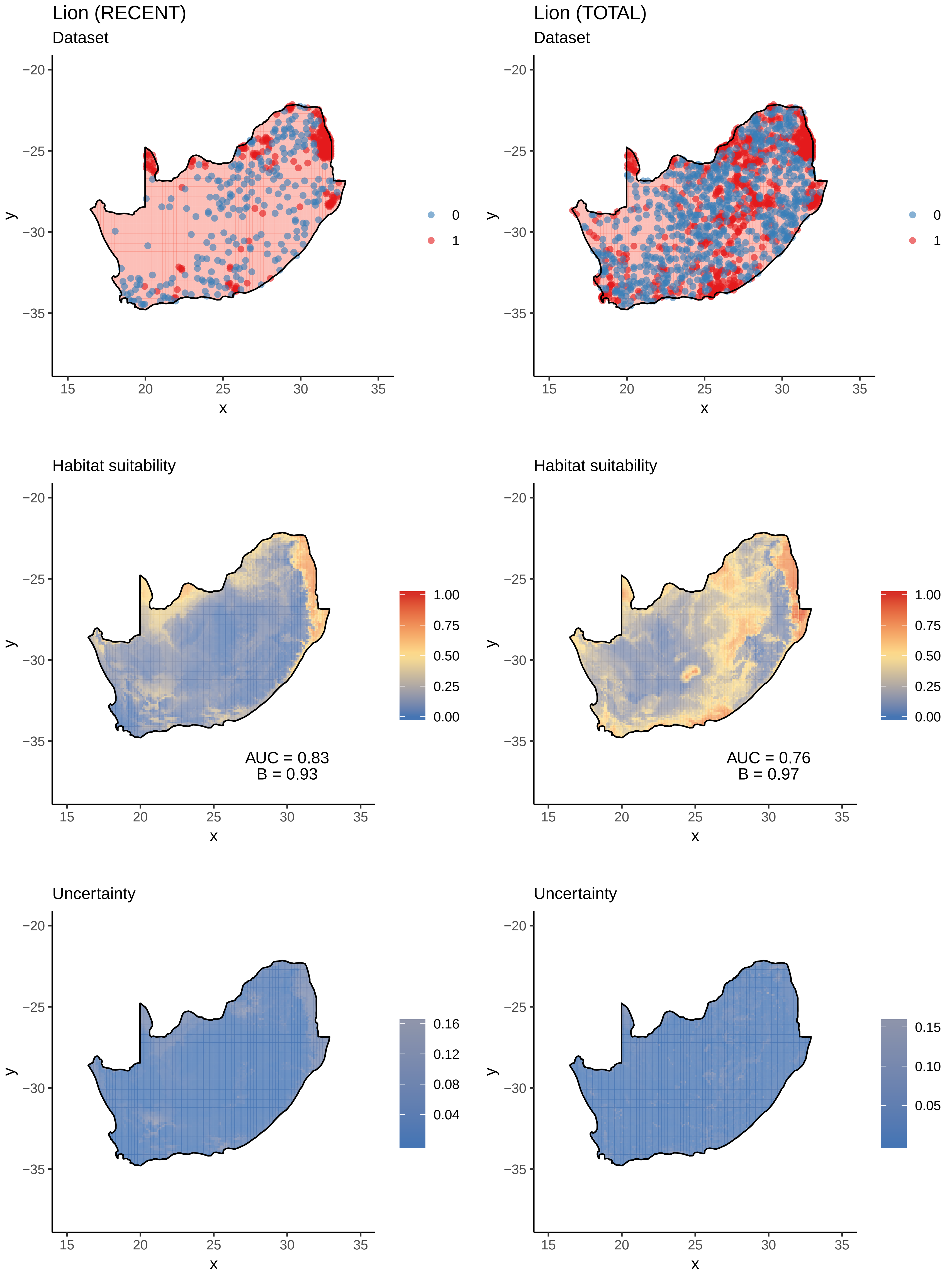

Lion (RECENT)
Dataset
Lion (TOTAL)
Dataset
0
1
Habitat suitability
AUC = 0.83
B = 0.93
AUC = 0.76
B = 0.97
1.00
0.75
0.50
0.25
0.00
Uncertainty
0.16
0.12
0.08
0.04
0.15
0.10
0.05
x
y

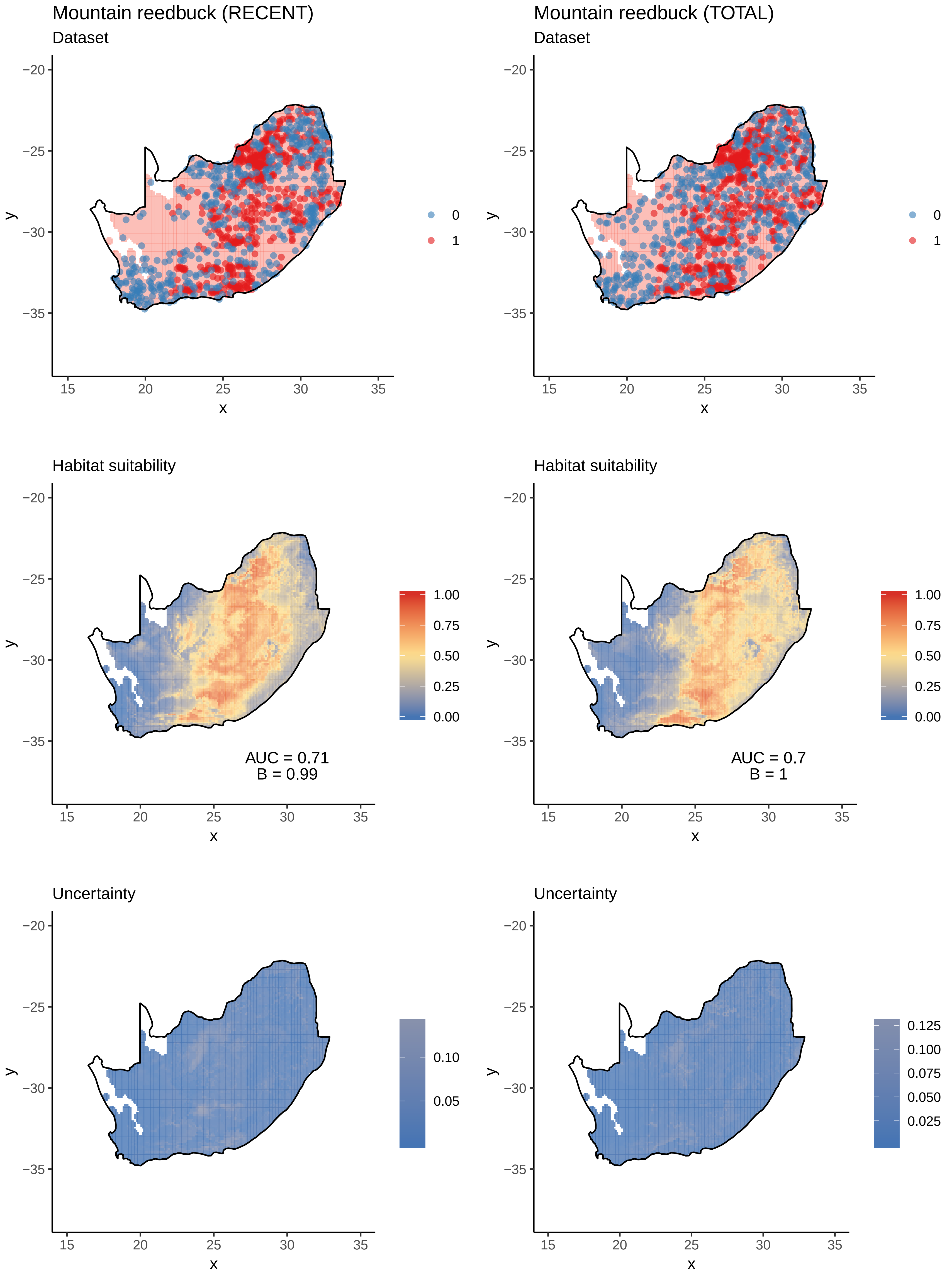

Mountain reedbuck (RECENT)
Dataset
Mountain reedbuck (TOTAL)
Dataset
0
1
y
x
−20
−25
−30
−35
15
20
25
30
35
Habitat suitability
1.00
0.75
0.50
0.25
0.00
AUC = 0.71
B = 0.99
AUC = 0.7
B = 1
Uncertainty
0.10
0.05
0.125
0.100
0.075
0.050
0.025

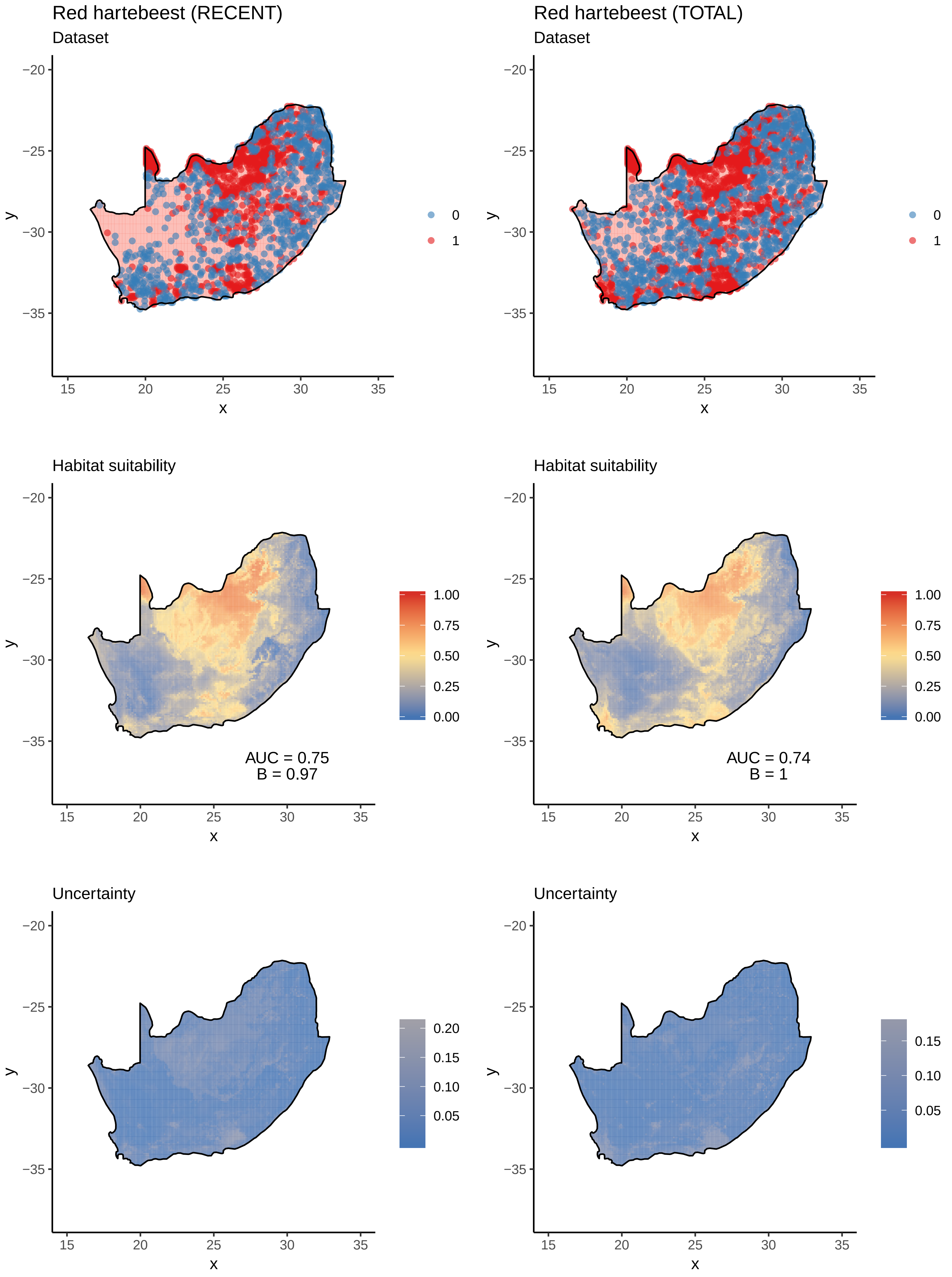

Red hartebeest (RECENT)
Dataset
Red hartebeest (TOTAL)
Dataset
Habitat suitability
AUC = 0.75
B = 0.97
Habitat suitability
AUC = 0.74
B = 1
Uncertainty
Uncertainty
x
y
0
1
1.00
0.75
0.50
0.25
0.00
0.20
0.15
0.10
0.05

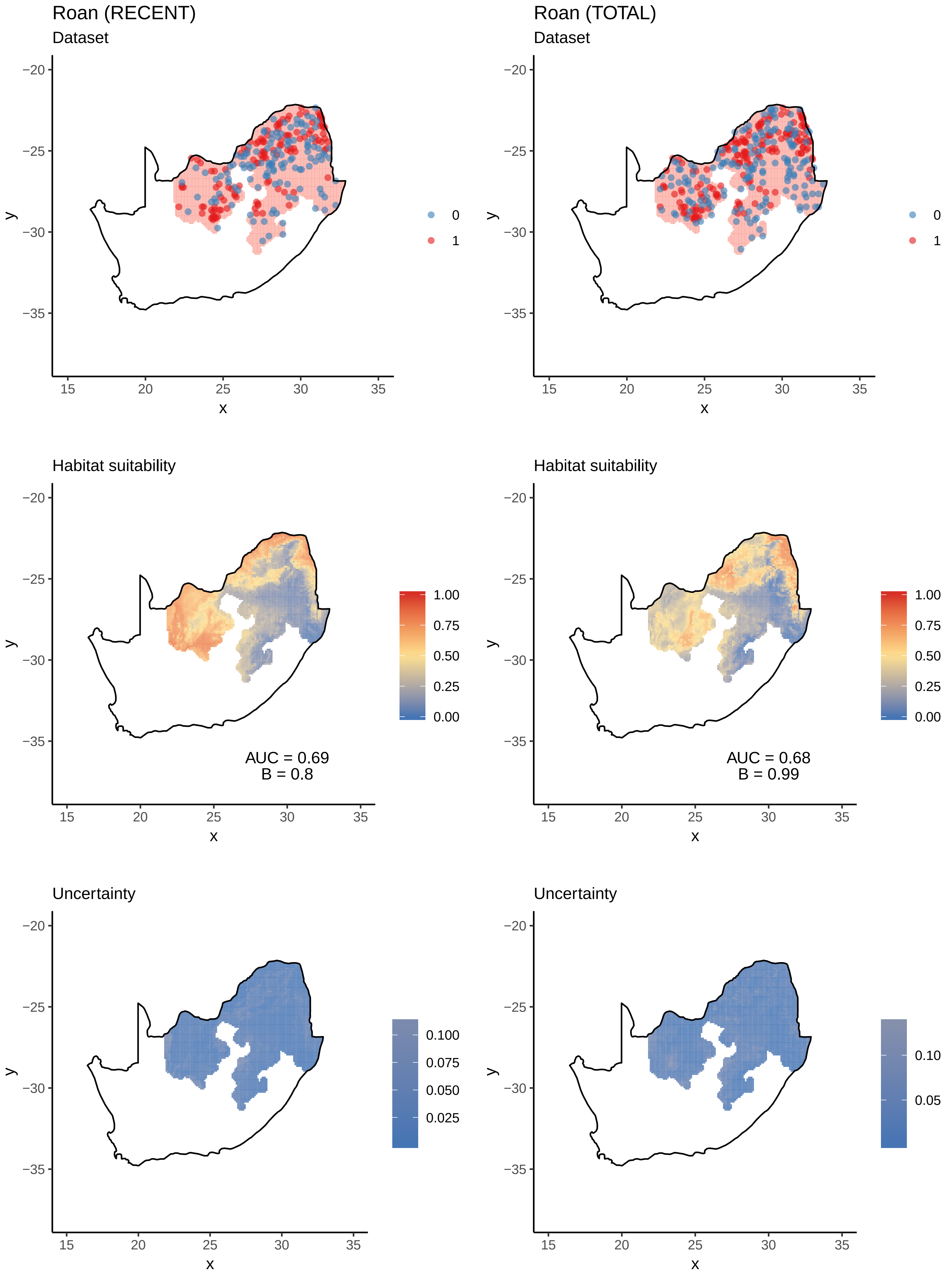
Roan (RECENT)
Dataset
Roan (TOTAL)
Dataset
0
1
Habitat suitability
AUC = 0.69
B = 0.8
Habitat suitability
AUC = 0.68
B = 0.99
1.00
0.75
0.50
0.25
0.00
Uncertainty
0.100
0.075
0.050
0.025
Uncertainty
0.10
0.05
x
y
−20
−25
−30
−35
15
20
25
30
35

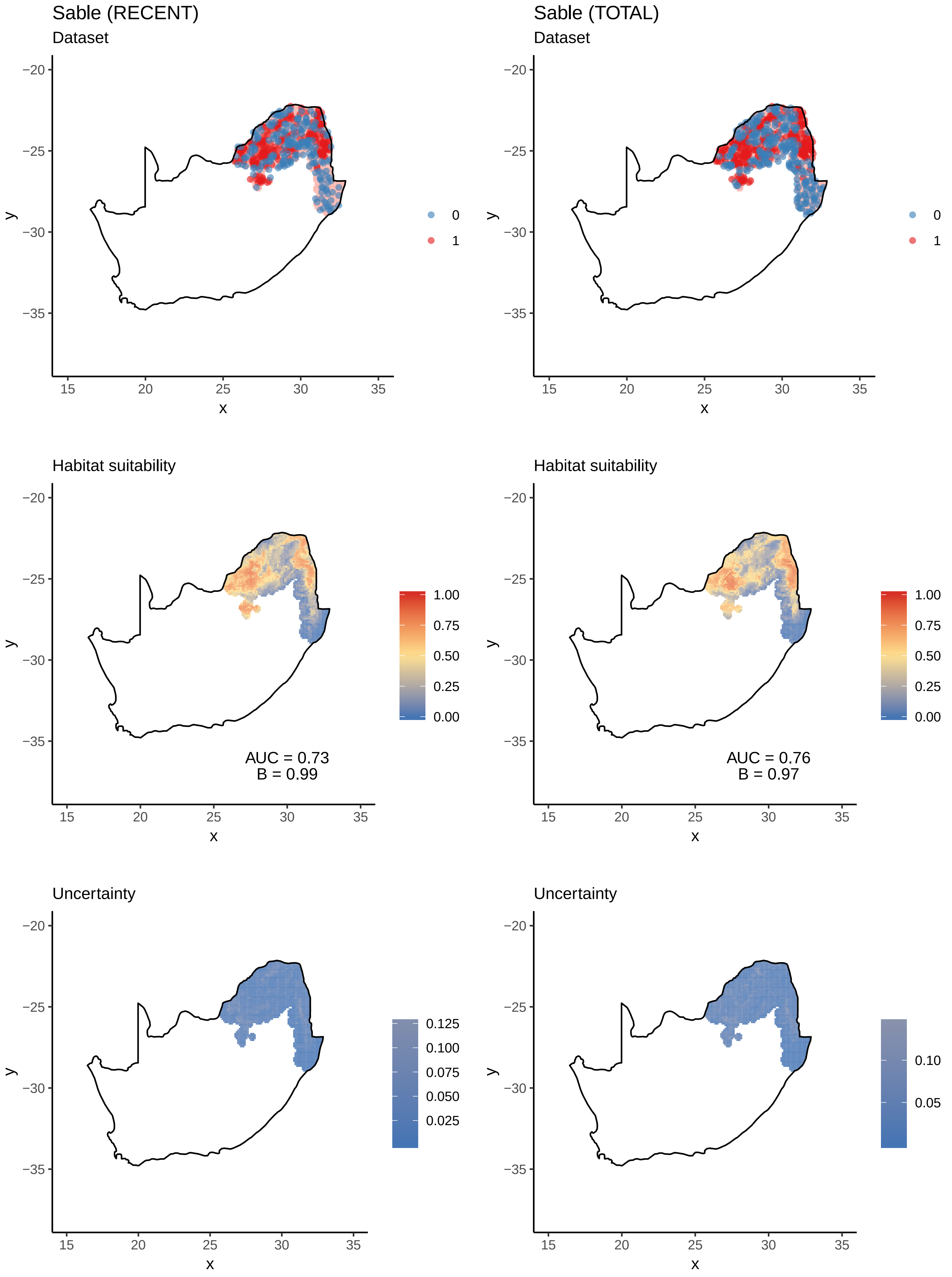

Sable (RECENT)
Dataset
Sable (TOTAL)
Dataset
Habitat suitability
AUC = 0.73
B = 0.99
Habitat suitability
AUC = 0.76
B = 0.97
Uncertainty
Uncertainty
x
y
−20
−25
−30
−35
15
20
25
30
35
0
1
1.00
0.75
0.50
0.25
0.00
0.125
0.100
0.075
0.050
0.025
0.10
0.05

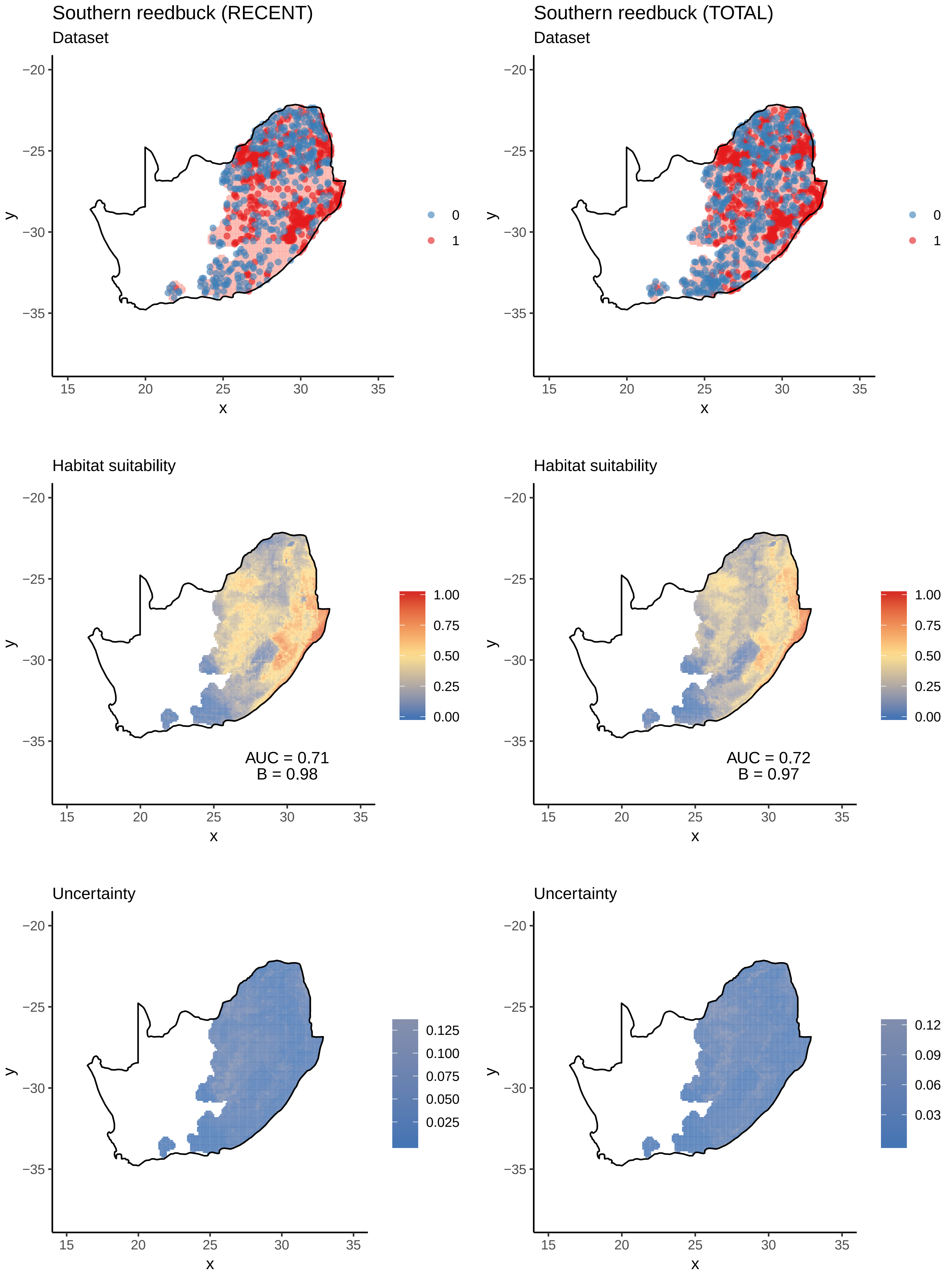
Southern reedbuck (RECENT)
Dataset
Southern reedbuck (TOTAL)
Dataset
0
1
Habitat suitability
1.00
0.75
0.50
0.25
0.00
AUC = 0.71
B = 0.98
AUC = 0.72
B = 0.97
Uncertainty
0.125
0.100
0.075
0.050
0.025
0.12
0.09
0.06
0.03
−20
−25
−30
−35
15
20
25
30
35
x
y

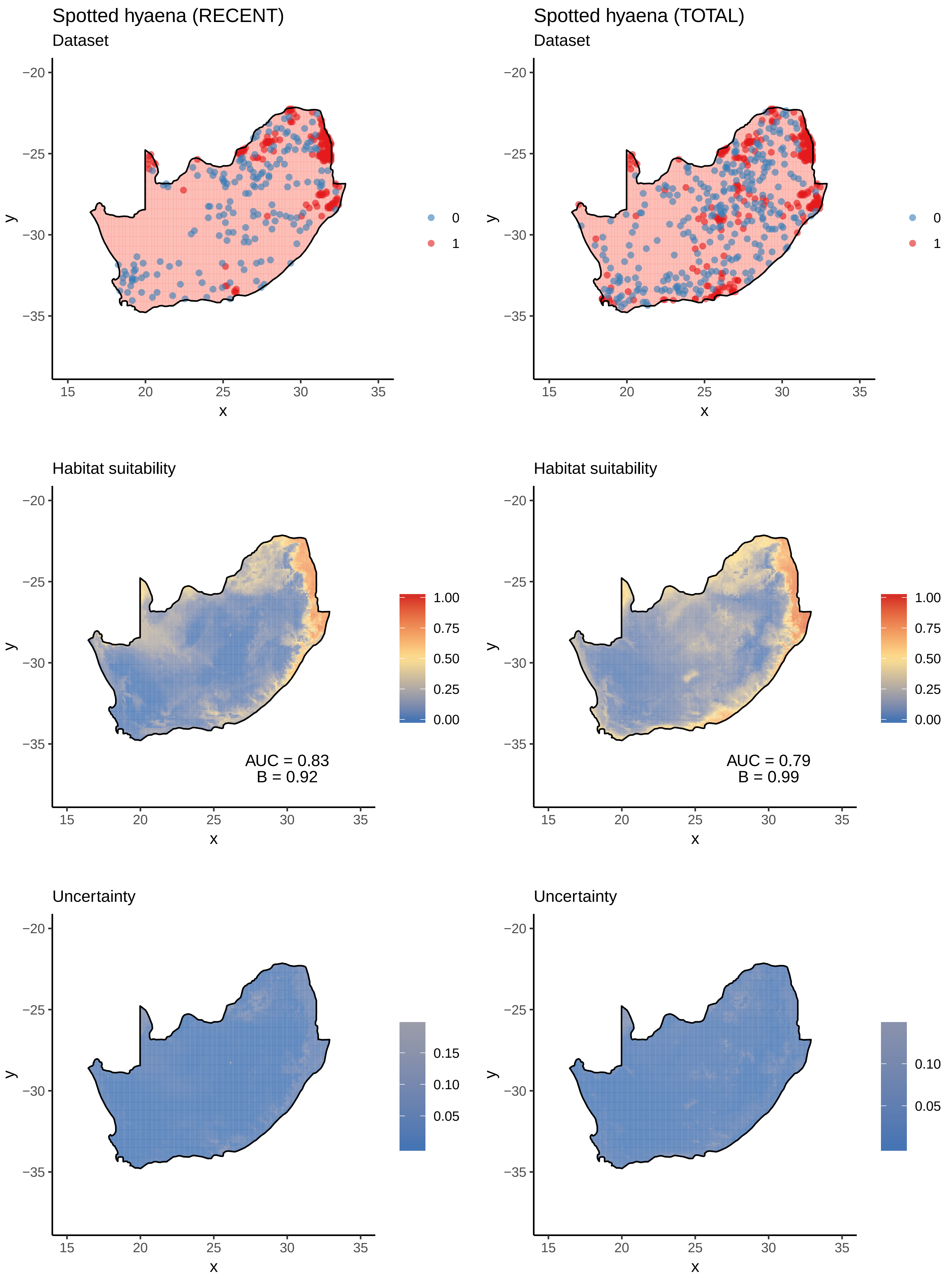

Spotted hyaena (RECENT)
Dataset
Spotted hyaena (TOTAL)
Dataset
0
1
Habitat suitability
AUC = 0.83
B = 0.92
AUC = 0.79
B = 0.99
1.00
0.75
0.50
0.25
0.00
Uncertainty
0.15
0.10
0.05
x
y

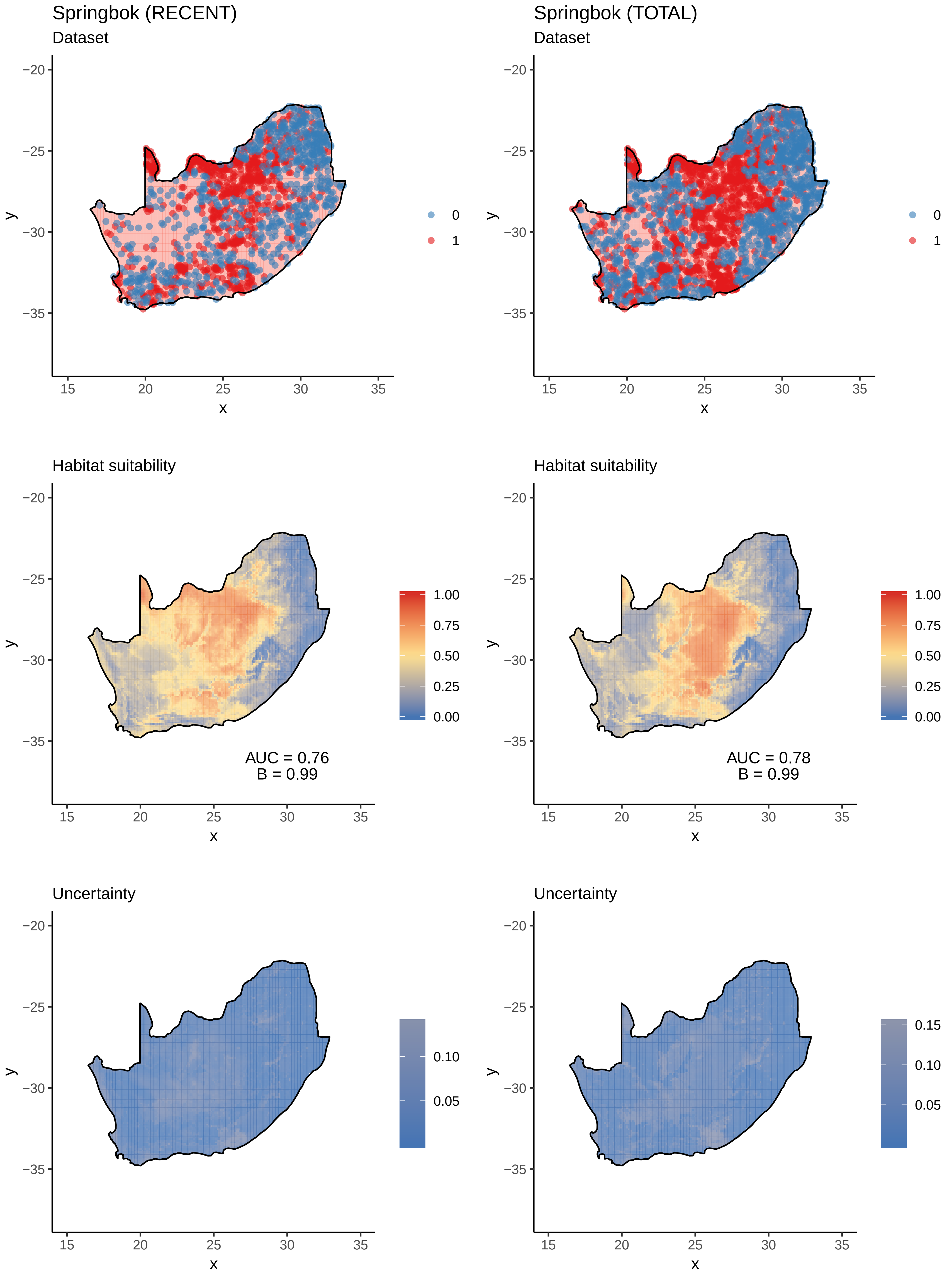

Springbok (RECENT)
Dataset
Springbok (TOTAL)
Dataset
0
1
Habitat suitability
AUC = 0.76
B = 0.99
AUC = 0.78
B = 0.99
1.00
0.75
0.50
0.25
0.00
Uncertainty
0.10
0.05
0.15
x
y

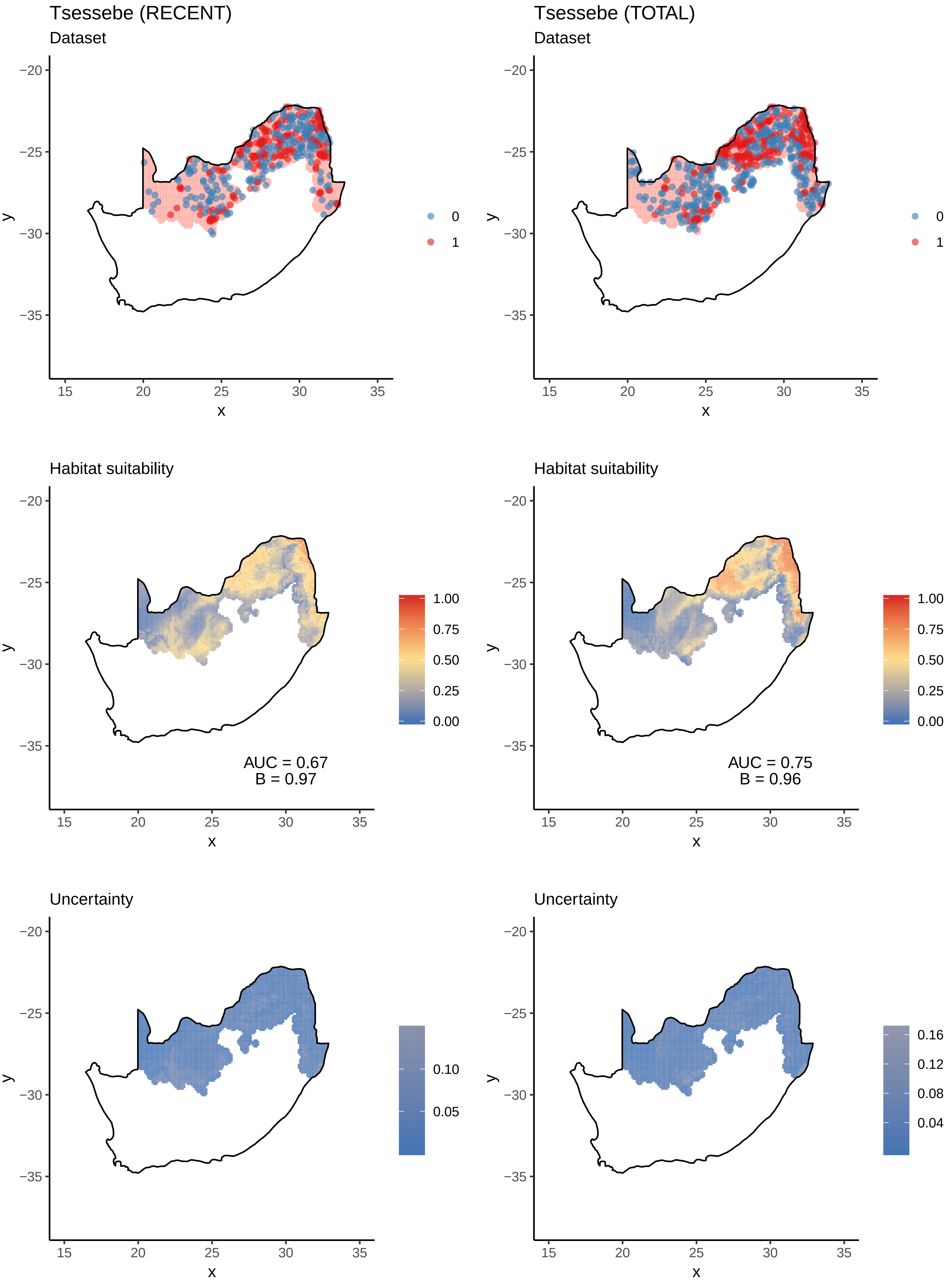
Tsessebe (RECENT)
Dataset
Tsessebe (TOTAL)
Dataset
Habitat suitability
AUC = 0.67
B = 0.97
AUC = 0.75
B = 0.96
Uncertainty
0
1
1.00
0.75
0.50
0.25
0.00
0.10
0.05
0.16
0.12
0.08
0.04
−20
−25
−30
−35
15
20
25
30
35
x
y

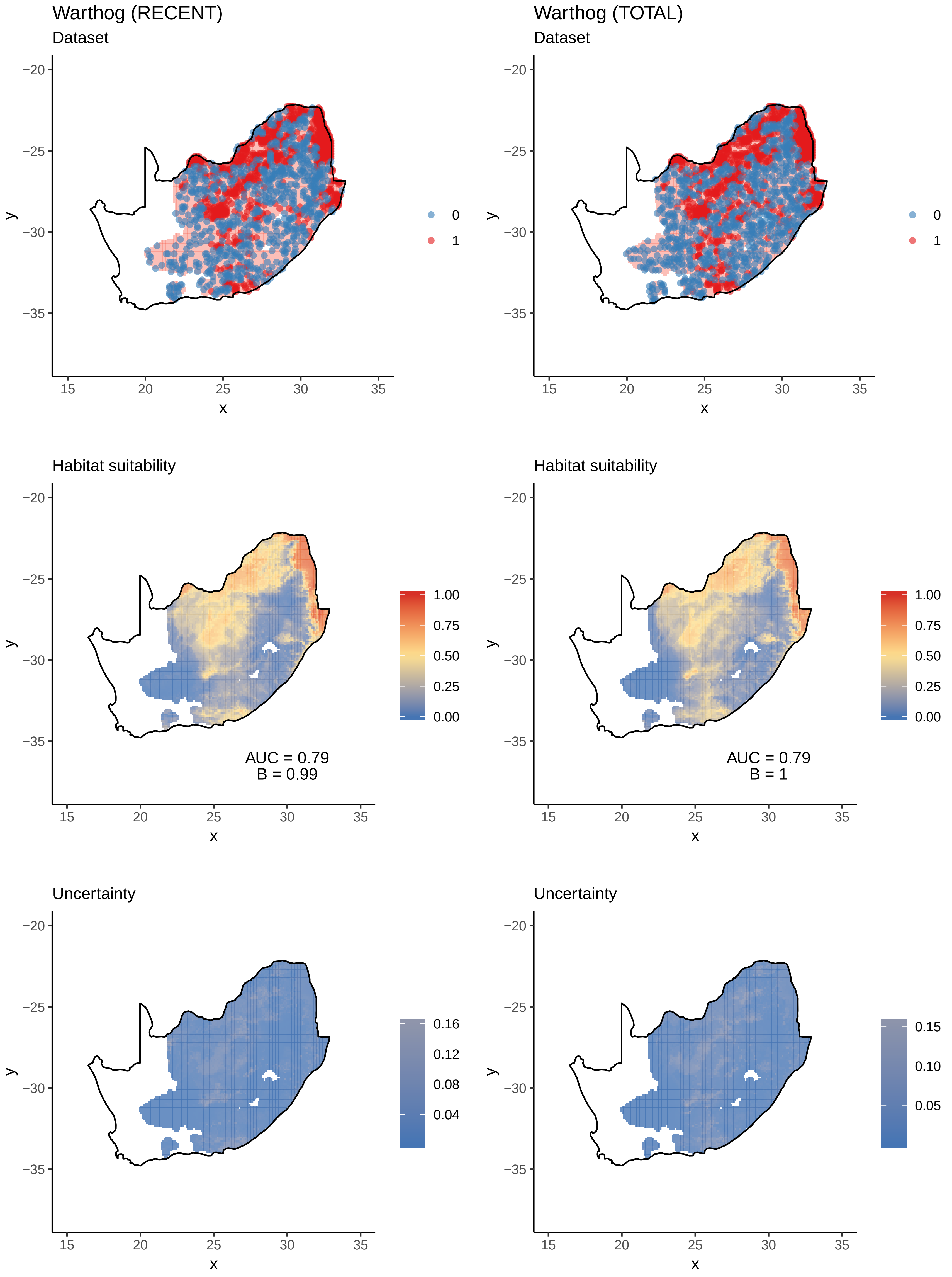

Warthog (RECENT)
Dataset
Warthog (TOTAL)
Dataset
0
1
Habitat suitability
AUC = 0.79
B = 0.99
AUC = 0.79
B = 1
1.00
0.75
0.50
0.25
0.00
Uncertainty
0.16
0.12
0.08
0.04
0.15
0.10
0.05
x
y
15
20
25
30
35
−20
−25
−30
−35

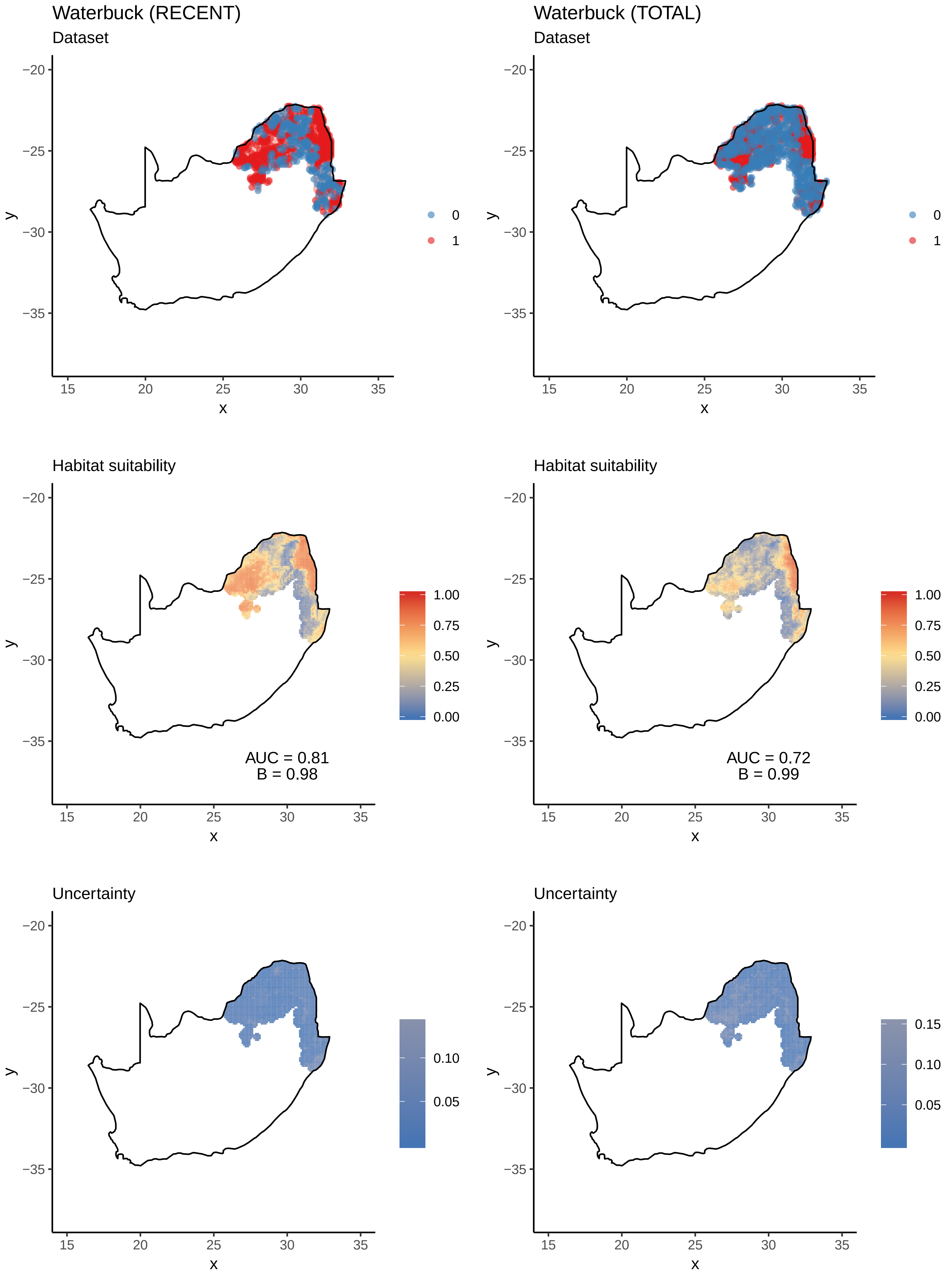

Waterbuck (RECENT)
Dataset
−20
−25
−30
−35
y
15
20
25
30
35
x
0
1
Waterbuck (TOTAL)
Dataset
0
1
Habitat suitability
1.00
0.75
0.50
0.25
0.00
AUC = 0.81
B = 0.98
Habitat suitability
AUC = 0.72
B = 0.99
Uncertainty
0.10
0.05
Uncertainty
0.15
0.10
0.05
